# Tumor-derived SAA1-TLR4 signaling drives tumor-to-muscle communication in pancreatic cancer cachexia

**DOI:** 10.64898/2026.09.04.748998

**Authors:** Gabriele Guarnaccia, Jimmy Massenet, Alessandra Cecchini, Elena Guarnaccia, Beatrice Silvestri, Chiara Nicoletti, Jesus R. Barajas, David Sala, Marcos G. Teneche, Brightany Li, Luca Caputo, Cedomir Stamenkovic, Tatiana M. Moreno, Rabi Murad, Shawn Delaware, Daphne Mayer, Yijuan Zhang, Priyanka Gupta, Alexandre R. Colas, Alessandro Vasciaveo, Caroline Kumsta, Peter D. Adams, Will Wang, Cesare Gargioli, Commisso Cosimo, Pier Lorenzo Puri, Alessandra Sacco

## Abstract

Cancer cachexia limits treatment tolerance and survival in pancreatic ductal adenocarcinoma (PDAC), yet the tumor-derived signals driving tissue dysfunction remain poorly understood. Here, we identify serum amyloid A1 (SAA1) as a mediator of tumor-to-host communication acting through Toll-like receptor 4 (TLR4). Tumor-derived SAA1 was elevated in human PDAC and in a mouse PDAC model and disrupted both myofiber and muscle stem cell (MuSC) homeostasis. Genetic reduction of tumor-derived SAA1 uncoupled tumor progression from host wasting, preserving muscle mass and function and prolonging survival without affecting primary tumor growth. Mechanistically, SAA1-TLR4 signaling drove multicellular remodeling of the skeletal muscle microenvironment. Therapeutic TLR4 inhibition after cachexia onset restored muscle mass, function and MuSC abundance and prolonged survival independently of tumor growth. Conservation of SAA1-TLR4 signaling in human skeletal muscle identifies a therapeutically actionable tumor–host pathway and demonstrates that host deterioration can be targeted independently of tumor progression.

## INTRODUCTION

Cancer cachexia is a multifactorial syndrome characterized by progressive loss of skeletal muscle and adipose tissue, systemic inflammation and metabolic dysfunction. Affecting up to 70% of patients with certain malignancies, cachexia reduces tolerance to anticancer therapies, diminishes quality of life, and is associated with poor survival^1,2^. Because cachexia cannot be reversed by conventional nutritional interventions^3^, identifying the mechanisms through which tumors drive dysfunction of distant tissues remains a major clinical challenge.

Pancreatic ductal adenocarcinoma (PDAC) has one of the highest incidences of cachexia, affecting more than 70% of patients ^4,5^. Skeletal muscle dysfunction is a major determinant of cachexia-associated morbidity and contributes to poor outcomes by compromising physical function and tolerance to anticancer therapy ^6^. Although inflammatory cytokines, including TNF-α, IL6, and IL-1β, contribute to skeletal muscle catabolism during cachexia, therapeutic strategies targeting cachexia have shown limited clinical benefit. Moreover, cachexia involves not only loss of muscle mass but disruption of tissue function and homeostasis, suggesting that mechanisms beyond canonical myofiber catabolism contribute to disease progression.

A prominent systemic response to inflammation is the acute phase response (APR), an evolutionarily conserved program characterized by production of circulating acute phase proteins ^7^. Although these proteins are consistently elevated in patients with cachexia, whether they are biomarkers of systemic inflammation or active mediators of tissue dysfunction remains unclear. Serum amyloid A1 (SAA1) is a highly inducible APR protein that is elevated in multiple malignancies and has been associated with muscle wasting in chronic inflammatory disorders ^8,9^. However, whether tumor-derived SAA1 acts as an endocrine mediator of skeletal muscle dysfunction during cachexia is unknown.

Here, we identify a tumor-to-host signaling axis in which pancreatic tumor-derived SAA1 activates TLR4 signaling to coordinately disrupt myofiber and muscle stem cell (MuSC) homeostasis. Integrating human patient data with orthotopic PDAC models, single-nucleus and spatial profiling, and human skeletal muscle systems, we establish SAA1-TLR4 signaling as a driver of multicellular muscle dysfunction. Genetic reduction of tumor-derived SAA1 or pharmacological inhibition of TLR4 uncouples tumor progression from host wasting, preserving skeletal muscle and extending survival without affecting primary tumor growth. These findings establish a previously unrecognized mechanism of tumor-host communication that redefines cancer cachexia as a multicellular failure of skeletal muscle homeostasis and identifies the SAA1-TLR4 signaling axis as a therapeutically actionable vulnerability.

## RESULTS

### Integrated human transcriptomic analyses identify SAA1 as a candidate mediator of tumor-host communication

To identify candidate mediators of tumor-host communication in PDAC, we integrated bulk RNA-sequencing datasets from human PDAC tumors (N=456) ^10^ and skeletal muscles from PDAC patients (rectus abdominis muscle, N=84) ^11^ derived from independent published cohorts. Comparative analysis identified 93 pathways commonly dysregulated in both tumor and skeletal muscle (**Figure 1A**), predominately involving inflammation (40%) and extracellular matrix (ECM) remodeling (25%), with additional tumor microenvironment/tissue damage (15%), coagulation (10%), and growth factor signaling/stress responses (10%) (**Figure 1B**).

**Figure 1.**
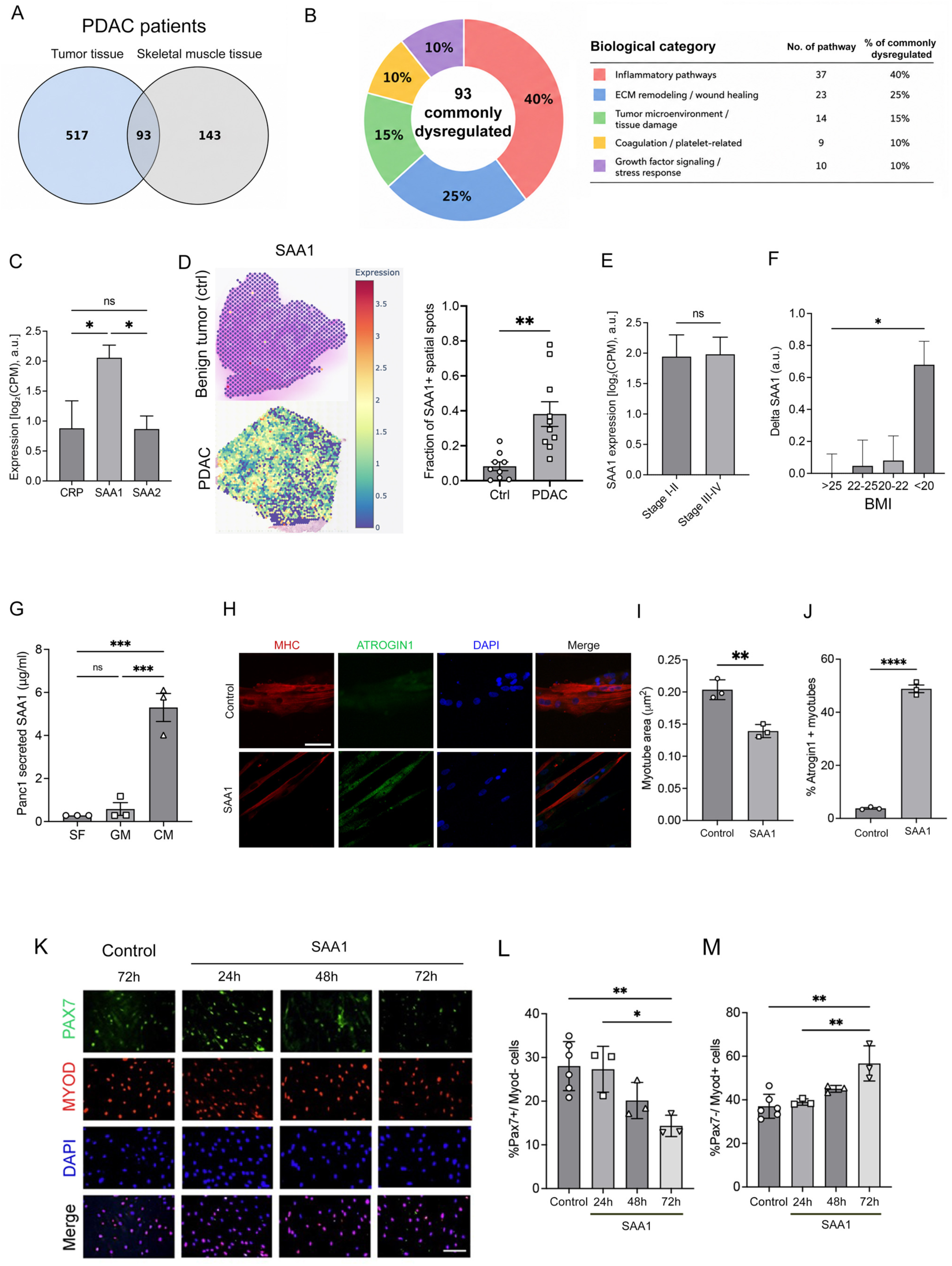
SAA-1 expression is elevated in human pancreatic ductal adenocarcinoma (PDAC) and induces human myotube atrophy and muscle progenitors’ myogenic commitment. (A) Venn diagram showing the overlap between pathways significantly dysregulated in cachexia within PDAC tumor tissue from the Bailey cohort (n = 456 patients) and skeletal muscle from the Bhatt cohort (n = 84 patients). In tumor tissue, 610 pathways were dysregulated, including 581 upregulated and 29 downregulated pathways. In skeletal muscle, 236 pathways were dysregulated, including 210 upregulated and 26 downregulated pathways. Ninety-three pathways were commonly dysregulated across both tissues. (B) Functional categorization of the 93 pathways shared between tumor and skeletal muscle according to their biological processes. Inflammatory pathways represented the largest category (40% of all shared pathways). Among these, five cytokine-related pathways were commonly upregulated in both tissues. (C) Expression levels of acute phase response genes (CRP, SAA1, and SAA2) in the Bailey et al PDAC tumor dataset ^10^. Data are shown as log2(CPM) normalized to non-tumor controls from UCSC Xena. Control = 0. (D) Spatial transcriptomic analysis of SAA1 expression in tumor tissue from patients with PDAC (n = 10) and non-tumoral controls with benign Pancreatic Intraductal Papillary-Mucinous Neoplasm (n = 9), using the DeepSpaceDB (visium 10x) ^12^. Quantification of the fraction of SAA1-positive spatial spots through Seurat. (E) Quantification of SAA1 expression by tumor stage (early vs. late) in the Bailey et al PDAC dataset ^10^. Expression values are reported as log2(CPM) normalized to healthy pancreas from UCSC Xena. N= 362 for UCSC Xena, N= 456 for PDAC. Control = 0. (F) Proteomic analysis from Cao et al ^14^ (n = 140 pancreatic cancer patients) showing the correlation between tumor SAA1 protein expression and body mass index (BMI). Data are normalized to the mean SAA1 expression of patients with BMI > 25, with outliers removed. Values are expressed as ΔSAA1 (arbitrary units). (G) Quantification of secreted SAA by ELISA in conditioned media from human pancreatic cancer cells (PANC-1), compared to control media and serum-free conditions. N=3. (H) Representative immunofluorescence images of MF20 (myosin heavy chain) and Atrogin-1 in NCAM-positive human-derived myotubes treated with recombinant SAA1 (10*μ*g/ ml) or vehicle control (MF20, red; Atrogin-1, green; DAPI, blue). Scale bar, 50*μ*m. (I) Quantification of MF20 fluorescence area in the presence or absence of recombinant SAA1. N= 3. (J) Quantification of the percentage of Atrogin-1+ myotubes in the presence or absence of recombinant SAA1 (10 *μ*g/ ml). N=3. (K) Representative immunofluorescence images of NCAM-positive human myogenic progenitors-derived cultures in growth conditions in the presence or absence of recombinant SAA1 (10 *μ*g/ ml) at the indicated time points (Pax7, green; MyoD, red; DAPI, blue). Scale bar, 100*μ*m. (L) Quantification of the percentage of Pax7+/MyoD-cells in the indicated conditions. N=3-6. (M) Quantification of the percentage of Pax7-/MyoD+ cells in the indicated conditions. N=3-6. For bioinformatic analyses, statistical significance was determined using methods appropriate for each analysis, including differential expression testing with multiple testing correction (Benjamini–Hochberg adjusted p-values). Only results with adjusted p < 0.05 are shown. Statistical analyses were performed using unpaired two-tailed Student’s t-test or one-way ANOVA, as appropriate. *p < 0.05, **p < 0.01, ***p < 0.005, and ****p < 0.001. Data are presented as mean ± SEM.

Among inflammatory programs enriched in both tissues, we prioritized the acute phase response (APR), which generates circulating proteins capable of signaling to distant tissues (**Suppl. Figure 1A-B**). *SAA1* was the most strongly upregulated APR-associated transcript in human PDAC tumors (**Figure 1C** and **Suppl. Figure 2A**). Spatial transcriptomic analysis of independent human PDAC datasets (10X Genomics Visium datasets in DeepSpaceBD ^12^) confirmed marked elevation of *SAA1* expression in tumor tissue relative to benign pancreatic lesions from age-matched controls (**Figure 1D**). *SAA2* expression was also increased but was substantially less abundant, whereas *CRP* was unchanged (**Suppl. Figure 1C-E**).

To define the temporal dynamics of *SAA1* expression during disease progression, we analyzed PDAC patient cohorts across tumor stages using cBioportal ^10^. *SAA1* was elevated from Stage I-II disease and remained high through stages III-IV (**Figure 1E** and **Suppl. Figure 2B**), while *SAA2* and *CRP* exhibited a milder upregulation (**Suppl. Figure 1F-G** and **Suppl. Figure 2C-D**). In contrast, *IL6* peaked during early-stage disease and later declined, while *TNF* did not show sustained elevated expression (**Suppl. Figure 1H-I** and **Suppl. Figure 2E-F**), suggesting that sustained *SAA1* production may represent a hallmark of PDAC associated systemic inflammation. Consistent with these findings, interrogation of scRNAseq from a human PDAC cohort further localized *SAA1* predominantly to tumor epithelial cells (**Suppl. Figure 1J**) ^13^.

PDAC tumor SAA1 protein abundance negatively correlated with body mass index (BMI) in an independent PDAC proteomic cohort (**Figure 1F** and **Suppl. Figure 2G**) ^14^.

In skeletal muscle (*rectus abdominis*) from patients with PDAC, SAA1 expression was also elevated in samples with a molecular atrophy signature defined by high expression of both atrogenes TRIM63 (MuRF1) and FBXO32 (Atrogin-1) ^11^ (**Suppl. Figure 1K**), linking SAA1 to muscle wasting in patients.

Consistent with a tumor source, human Panc1 PDAC cells ^15^ secreted high levels of SAA1 into conditioned media (CM), whereas IL6 secretion was not significantly increased (**Figure 1G** and **Suppl. Figure 1L**). We next investigated whether SAA1 directly affects skeletal muscle cells. Recombinant SAA1 reduced myotube area and increased Atrogin-1 in primary human myotubes ^16 17^ (**Figure 1H-J**). Given the emerging role of MuSC dysfunction in cancer cachexia ^18–22^, we additionally examined the effect of SAA1 on human myogenic progenitors. SAA1 exposure promoted myogenic commitment, characterized by a progressive reduction in the percentage of Pax7+MyoD-cells, and an increase in the percentage of committed Pax7-MyoD+ cells (**Figure 1K-M**). Thus, SAA1 is produced by PDAC cells, associates with systemic and skeletal muscle wasting in patients, and directly disrupts human myofiber and MuSC homeostasis.

### Systemic SAA elevation and muscle dysfunction precede overt wasting in orthotopic PDAC

To define the temporal progression of PDAC cachexia *in vivo*, we orthotopically implanted syngeneic KPC cells (murine PDAC cells) into the pancreata of 8-10-week-old C57BL6 mice (**Figure 2A**) ^23–27^. Tumor-bearing mice developed substantial tumor burden and reduced tumor-free body weight (BW) by five weeks (**Figure 2B-C** and **Suppl. Figure 3A**). Tibialis anterior (TA) and gastrocnemius muscle weights normalized to BW declined beginning four weeks after KPC implantation, whereas soleus mass remained largely preserved (**Figure 2D-F** and **Suppl. Figure 3B-D**), consistent with the relative resistance of oxidative muscles to cachexia ^28–30^. TA myofiber cross-sectional area (CSA) similarly decreased from four weeks onward (**Figure 2G-H** and **Suppl. Figure 3E**). In contrast, soleus myofiber CSA was not significantly altered (**Suppl. Figure 3H-I**). Multiplex spatial imaging using CODEX ^31–33^ further revealed increased Atrogin-1 in cachectic TA myofibers relative to sham-treated controls (**Figure 2I-J**). No significant changes in fiber type composition were observed (**Suppl. Figure 3F-G**), indicating that the reduction in muscle mass primarily reflected myofiber atrophy rather than fiber-type switching.

**Figure 2.**
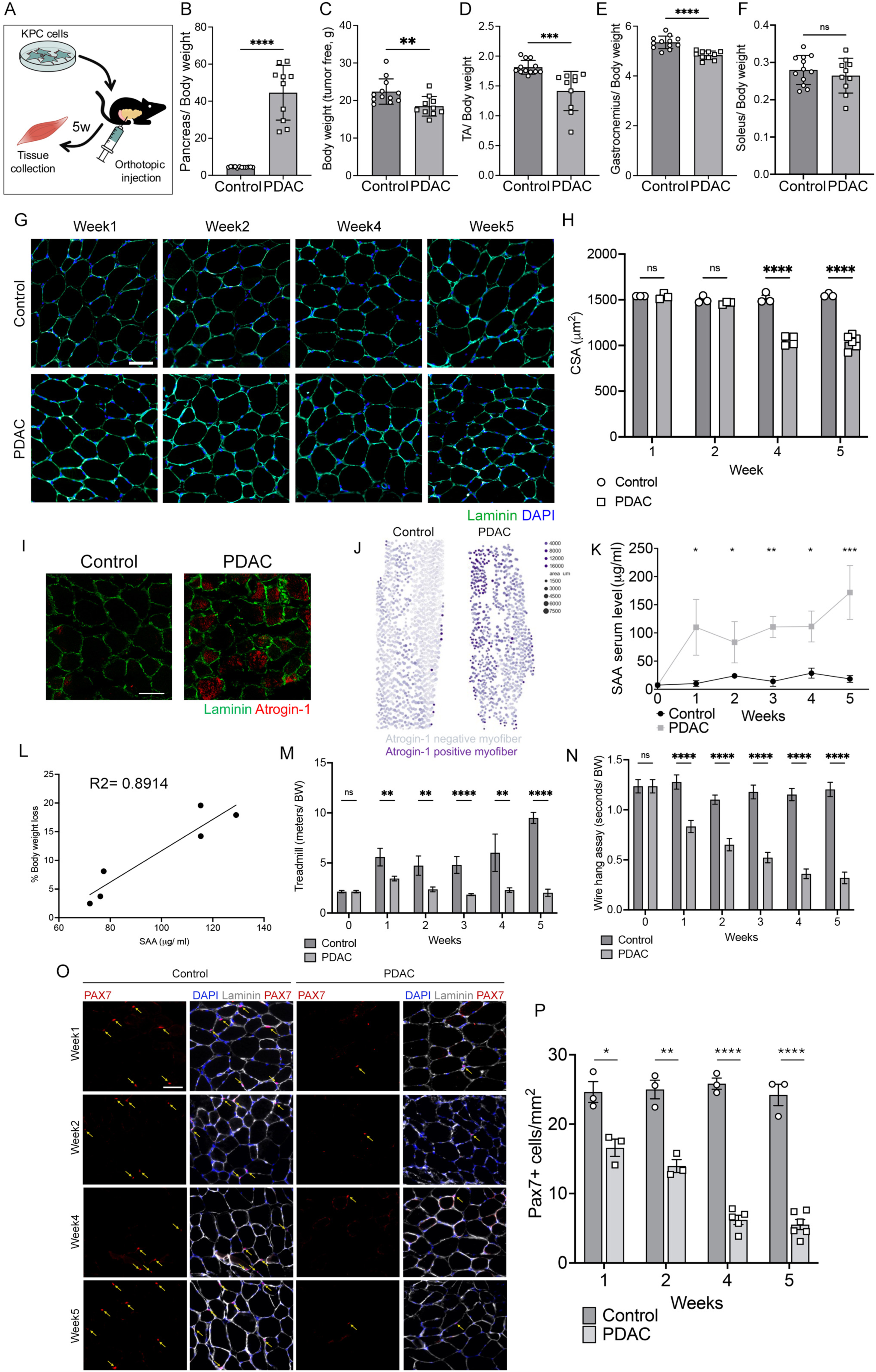
Orthotopic KPC cell injection recapitulates PDAC cachexia. (A) Schematic of the orthotopic transplantation of 25,000 KPC cells *in vivo*. (B) Quantification of pancreas weight normalized to body weight (mg/g) of mice transplanted with KPC cells or sham-treated controls at 5 weeks post cell injection. N= 10-12. (C) Quantification of tumor free body weight of mice transplanted with KPC cells or sham-treated controls at 5 weeks post cell injection. N= 10-12. (D) Quantification of weight of tibialis anterior muscle (TA), normalized to body weight of mice transplanted with KPC cells or sham-treated controls at 5 weeks post cell injection (mg/g). N= 10-12. (E) Quantification of weight of gastrocnemius muscle normalized to body weight of mice transplanted with KPC cells or sham-treated controls at 5 weeks post cell injection (mg/g). N= 10-12. (F) Quantification of weight of soleus muscles normalized to body weight of mice transplanted with KPC cells or sham-treated controls at 5 weeks post cell injection (mg/g). N= 10-12. (G) Representative immunofluorescence images of skeletal muscle cross-sections of TA muscle of mice transplanted with KPC cells or sham-treated controls at the indicated time points post tumor implantation (laminin, green; DAPI, blue). Scale bar, 100*μ*m. (H) Quantification of myofiber cross-sectional area (CSA) of TA muscles of mice transplanted with KPC cells or sham-treated controls at the indicated time points post tumor implantation. N= 3-6. (I) Representative images of CODEX immunofluorescence of TA muscle of mice transplanted with KPC cells or sham-treated controls at 5 weeks post cell injection (laminin, green; Atrogin-1, red). Scale bar, 100*μ*m. (J) Spatial quantification of Atrogin-1 expression at the single-myofiber level. Individual myofibers were computationally segmented and represented as dots, preserving their spatial organization within the muscle cross-section. Dot color intensity reflects Atrogin-1 expression levels, enabling visualization of the spatial distribution and heterogeneity of atrophic signaling across the tissue. Dot size represents myofibers CSA. N=3. (K) ELISA quantification of serum SAA levels in mice transplanted with KPC cells or sham-treated controls at the indicated time points post tumor implantation. N=3. (L) Correlation of serum SAA levels and percentage of tumor free body weight loss in individual KPC-injected mice at 5 weeks post tumor implantation. (M) Longitudinal treadmill performance assessment over 5 weeks in mice transplanted with KPC cells or sham-treated controls. Quantification of running distance (meters) measured weekly and normalized to body weight at each time point. N=6-27. (N) Longitudinal wire hang test assay over 5 weeks in mice transplanted with KPC cells or sham-treated controls. Hanging time (seconds) was recorded weekly and normalized to body weight at each time point. N=6-27. (O) Representative immunofluorescence images of TA skeletal muscle cross-sections of mice transplanted with KPC cells or sham-treated controls at the indicated time points post tumor implantation (laminin, white; Pax7, red; DAPI, blue). Scale bar, 100*μ*m. (P) Quantification of number of Pax7+ MuSC in TA skeletal muscle cross-sections of mice transplanted with KPC cells or sham-treated controls at the indicated time points post tumor implantation. N=3-6. Statistical analyses were performed using unpaired two-tailed Student’s t-test. Significance is indicated as follows: *p < 0.05, **p < 0.01, ***p < 0.005, and ****p < 0.001. Data are presented as mean ± SEM.

Circulating SAA increased within one week of tumor implantation, remained elevated throughout disease progression, and preceded detectable muscle wasting (**Figure 2K** and **Suppl. Figure 4A**). Serum proteomics at four weeks identified SAA1 as the predominant circulating isoform, with a modest increase in SAA2, in cachectic mice (**Suppl. Figure 3K**). Circulating SAA levels also positively correlated with tumor-free BW loss across individual animals (**Figure 2L**), supporting a close relationship between systemic SAA elevation and cachexia severity.

Notably, treadmill endurance and wire hang assay performance were impaired within one week of tumor implantation, before measurable reductions in muscle mass or myofiber CSA (**Figure 2M-N** and **Suppl. Figure 4B-C**). This early functional decline was accompanied by depletion of Pax7+ MuSC in TA muscle, whereas the soleus was largely unaffected (**Figure 2O-P** and **Suppl. Figure 3H,J**). Flow cytometry independently confirmed reduced MuSC abundance in cachectic muscle (**Suppl. Figure 3L**). Thus, systemic SAA elevation, muscle dysfunction and MuSC depletion precede overt myofiber atrophy during PDAC progression, revealing early disruption of skeletal muscle homeostasis before structural wasting becomes apparent.

### Single-nucleus transcriptomics reveals multicellular disruption of skeletal muscle homeostasis during cachexia

To define the cellular and transcriptional landscape of PDAC cachexia, we performed single nucleus RNAseq (snRNAseq) ^34–36^ of gastrocnemius muscles of orthotopic tumor-bearing or sham-treated mice five weeks after tumor implantation. After quality control, we analyzed 33,619 control and 32,129 cachectic nuclei, identifying 16 distinct meta-clusters corresponding to major muscle-resident cell types and myonuclear populations (**Figure 3A-B** and **Suppl. Figure 5A**). Cachectic muscles showed broad remodeling of cellular composition, including altered myonuclear states and reductions in MuSC, fibroadipogenic progenitors (FAP), and endothelial cells (**Figure 3A-C**). Pseudobulk analysis revealed a catabolic and metabolically altered transcriptional program (**Figure 3D** and **Suppl. Figure 5B**), marked by increased ubiquitin-proteasome and autophagy genes (*Trim63, Fbxo32, Bnip3, Gabarapl1, Retreg1*), and lipid metabolism genes (*Pdk4, Cd36, Pnpla2, Angptl4*), together with inflammatory and stress-response programs, including *Il6ra*, *Cebpd*, *Egln3*, and metallothioneins (*Mt1/2*). Conversely, extracellular matrix (ECM) (*Col1a1*, *Col1a2*, *Col3a1*, *Col6a2/3*, *Postn*, *Has2*, *Fstl1*), fast-twitch contractile components (*Myh4*, *Mybph*), and neuromuscular signaling genes (*Nrxn3*, *Robo1*, *Kcnh7*) were downregulated, consistent with broader disruption of structural, metabolic and homeostatic programs.

**Figure 3.**
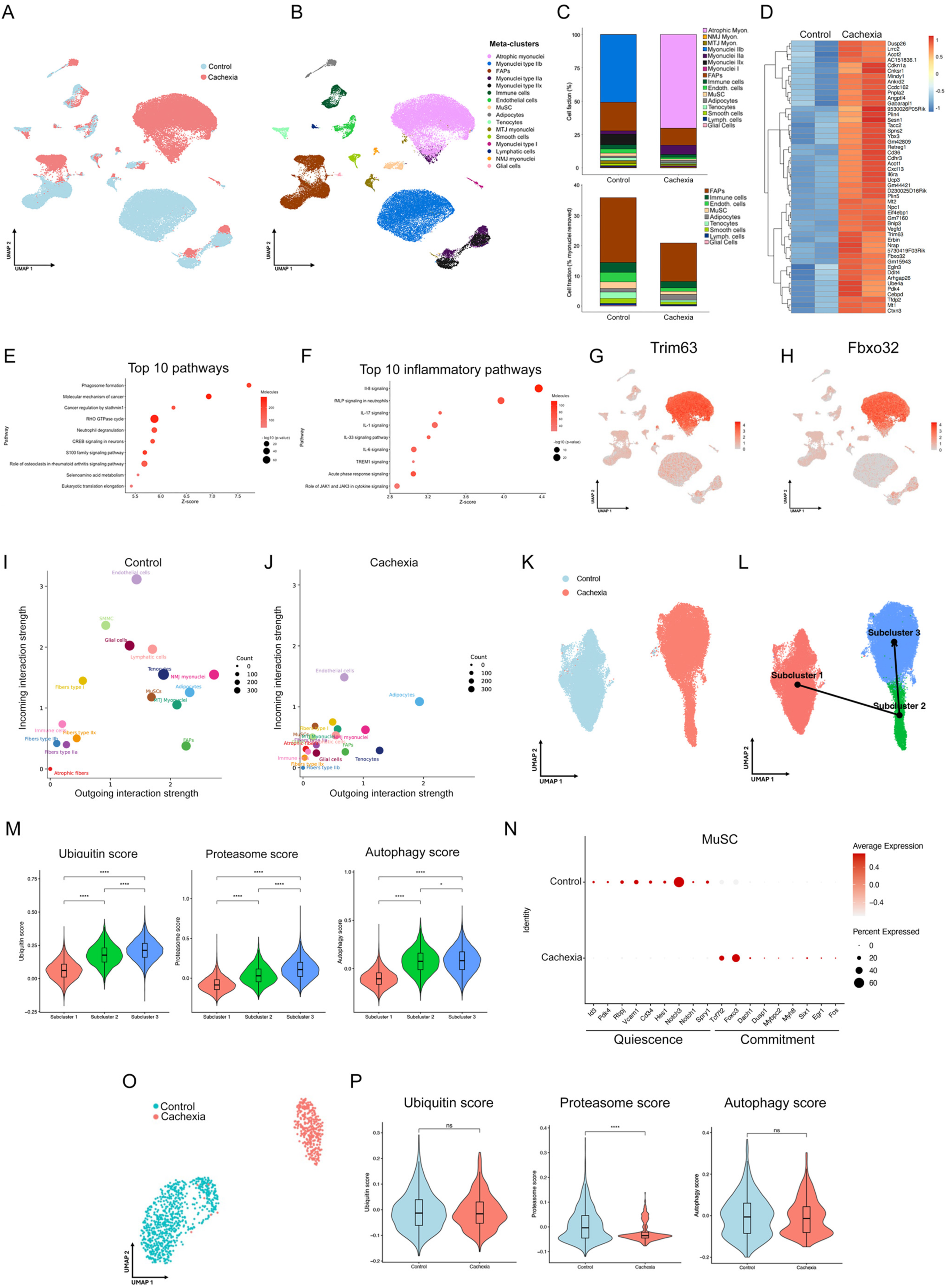
Single-nucleus transcriptomic characterization reveals progressive myonuclear atrophy and disruption of tissue homeostasis in cancer cachexia. (A) Umap visualization of single-nucleus-RNA-seq (snRNAseq) data of gastrocnemius muscle from KPC-injected or sham-treated controls at 5 weeks post tumor implantation. (B) Cell type annotation of identified meta-clusters of snRNAseq based on canonical marker genes. (C) Cell-type composition analysis showing the fraction of each indicated cell population. Top, all nuclei; bottom, all mononucleated cell types (excluding skeletal muscle myonuclei). (D) Top 50 upregulated genes in PDAC cachexia vs. sham-treated controls identified by pseudobulk differential expression (DEG) analysis across whole muscle. Key atrophy-related genes (Trim63, Fbxo32) and inflammation-associated genes (Mt1, Mt2, Il6ra) are shown. (E) Ingenuity Pathway Analysis (IPA) of whole-muscle transcriptomic changes in PDAC cachexia showing the top 10 enriched pathways, including muscle degradation-associated pathways (phagosome formation), inflammatory signaling (S100 family signaling), and cytoskeletal remodeling pathways (Rho GTPase signaling). (F) IPA of inflammation-related pathways in whole muscle highlighting enrichment of acute phase response signaling among the top 10 pathways in cachexia. (G-H) UMAP feature plots showing expression of Trim63 (MuRF1) (G) and Fbxo32 (Atrogin-1) (H) across all nuclei. (I-J) Cell-cell communication analysis using CellChat in sham-treated controls (I) and PDAC cachexia (J) conditions. (K) Subclustering of type IIb and atrophic myonuclei populations to resolve transcriptional heterogeneity. (L) Pseudotime trajectory analysis (Monocle3) illustrating the progression and transcriptional transition from type IIb to atrophic myonuclei states. (M) Module score analysis of ubiquitin and proteasome system and autophagy pathways across identified myonuclear subclusters, supporting progressive activation of proteolytic programs during atrophy. (N) Dot plot showing expression of genes associated with MuSC quiescence and myogenic commitment in MuSC from KPC-injected and sham-treated mice. Dot size indicates the fraction of cells within each group expressing the indicated gene (%), whereas color intensity represents the average expression level within each group. (O) Subclustering of MuSC populations to resolve transcriptional heterogeneity. (P) Module score analysis of ubiquitin and proteasome system and autophagy pathways across identified MuSC subclusters. For bioinformatic analyses, statistical significance was determined using methods appropriate for each analysis, including differential expression testing with multiple testing correction (Benjamini–Hochberg adjusted p-values). Only results with adjusted p < 0.05 are shown. Statistical analyses were performed using unpaired two-tailed Student’s t-test or one-way ANOVA, as appropriate. Significance is indicated as follows: *p < 0.05, ***p < 0.005. Data are presented as mean ± SEM.

Pathway analysis similarly identified activation of inflammatory programs, including S100, IL-17, acute phase response, TREM1 and JAK/STAT signaling, accompanied by suppression of metabolic and homeostatic pathways (**Figure 3E-F**). In contrast, downregulated pathways were enriched for metabolic and homeostatic programs, including branched-chain amino acid catabolism, glycogen metabolism, PPAR signaling, RHOGDI signaling, and mitophagy, consistent with impaired nutrient utilization, defective mitochondrial quality control, and loss of metabolic flexibility in cachectic muscle (**Suppl. Figure 5C**). *Trim63* and *Fbxo32* were enriched predominantly within atrophic myonuclear populations (**Figure 3G-H** and **Suppl. Figure 5A**). CellChat analysis^37^ further revealed a widespread reduction of intercellular communication throughout the cachectic muscle microenvironment (**Figure 3I-J** and **Suppl. Figure 6A-B**).

To resolve the progression of myonuclear remodeling, we reclustered type IIb and atrophic myonuclei ^37,38^, identifying three transcriptionally distinct subpopulations (**Figure 3K**). Trajectory inference using Monocle3 ^39^ positioned these subpopulations along a continuum from healthy to progressively cachectic states (**Figure 3L**), accompanied by increasing ubiquitin-proteasome and autophagy module scores (**Figure 3M**). Proteostatic remodeling extended across multiple stromal and non-myogenic populations (**Suppl. Figure 5D-F**), indicating that cachexia induces tissue-wide stress rather than an exclusively myofiber-autonomous response.

MuSC, however, underwent a distinct pathological transition. Unlike cachectic myonuclei, MuSC did not activate ubiquitin-proteasome or autophagy programs and instead showed reduced proteasome scores (**Figure 3O-P**). MuSC exhibited reduced expression of quiescence-associated genes and increased myogenic commitment programs (**Figure 3N**), consistent with premature activation and the progressive depletion observed *in vivo*. Their interactions with surrounding niche populations were also broadly diminished (**Suppl. Figure 5G-H**). Thus, PDAC cachexia induces distinct pathological states across muscle-resident populations, characterized by progressive proteostatic remodeling of myonuclei, loss of MuSC quiescence and widespread disruption of intercellular communication.

### Tumor-derived SAA1 drives skeletal muscle dysfunction during PDAC cachexia

To determine whether SAA1 is sufficient to reproduce the features of cachectic muscle dysfunction, we exposed freshly isolated MuSC from skeletal muscle of C57BL6 mice at 12-14 weeks of age to KPC conditioned medium (CM) or recombinant SAA1 in growth conditions for 48hrs. Both treatments reduced the proportion of Pax7+MyoD-cells and increased committed Pax7-MyoD+ cells and differentiating myogenin+ populations (**Suppl. Figure 5I-J**), consistent with the loss of MuSC quiescence observed *in vivo*. In differentiated primary myotubes, KPC CM and recombinant SAA1 similarly induced atrophy (**Figure 4A**), consistent with previous reports in C2C12 myoblasts ^40^. We next administered recombinant SAA1 intramuscularly into the TA muscles of tumor-free mice at 12-14 weeks of age every 2 days, and tissues harvested 6 days after treatment initiation (**Figure 4B**). SAA1 reduced myofiber CSA, and depleted Pax7+ MuSC relative to vehicle-treated controls (**Figure 4C-E**). Consistent with these findings, qPCR analysis revealed increased expression of the atrophy-related gene Atrogin-1 (*Fbxo32*) in both SAA1-injected muscles (**Suppl. Figure 5K**), and in PDAC cachectic muscle in mice relative to healthy controls (**Suppl. Figure 5L**), demonstrating that SAA1 is sufficient to reproduce key features of cachectic muscle remodeling *in vivo*.

**Figure 4.**
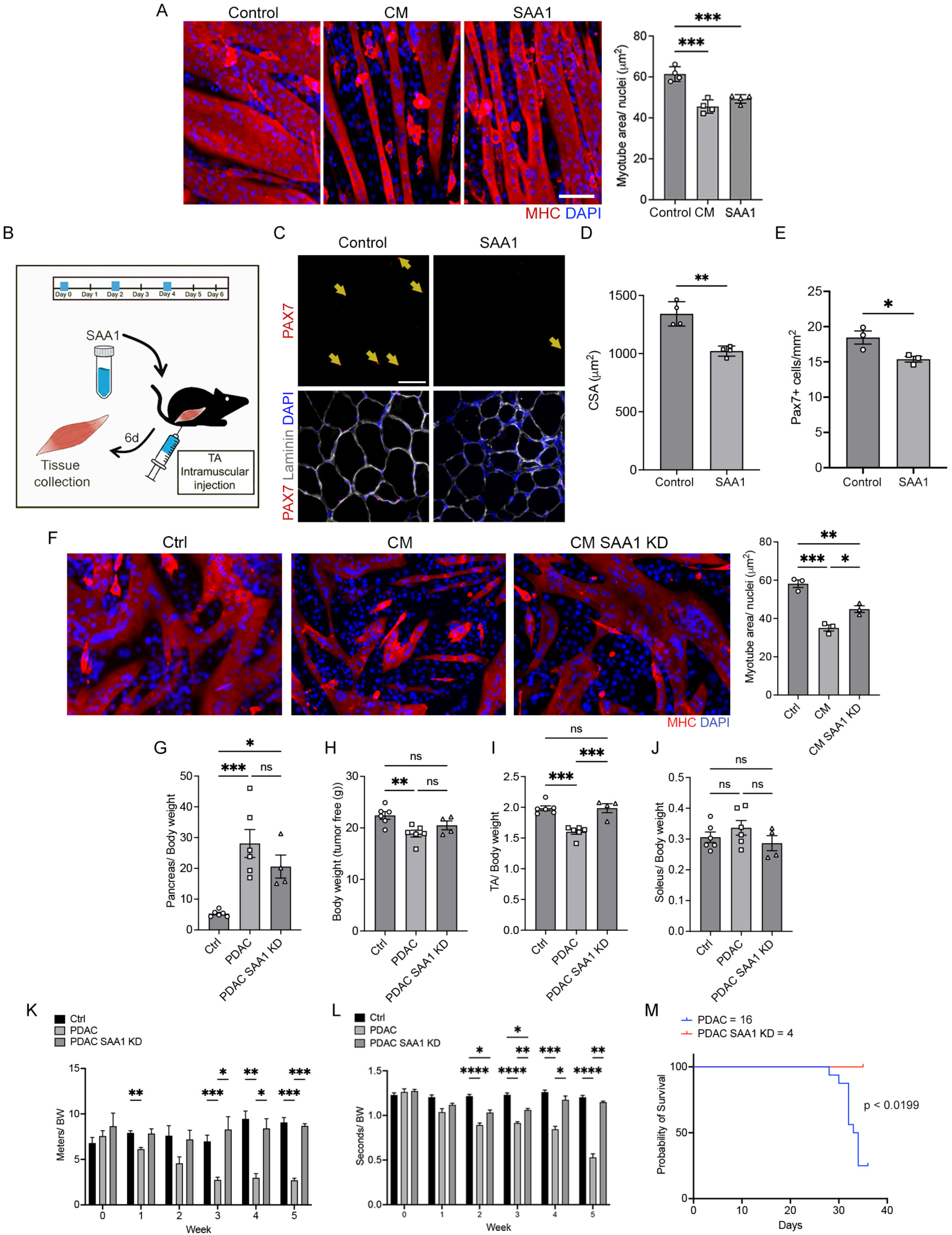
Depletion of tumor-derived SAA1 attenuates pancreatic cancer-induced muscle wasting and functional decline. (A) Representative immunofluorescence images of mouse MuSC-derived myotubes treated with KPC conditioned media (CM), recombinant SAA1 (10 *μ*g/ ml) or vehicle control for 48 hours (myosin heavy chain, MF20, red; DAPI, blue) (left). Scale bar, 50*μ*m. Quantification of myotube area normalized to the number of nuclei of mouse MuSC-derived myotubes treated with KPC conditioned media (CM), recombinant SAA1 (10 *μ*g/ ml) or vehicle control for 48 hours (right). N=4. (B) Schematic of treatment of wild-type C57BL6 mice by intramuscular injection of recombinant SAA-1. (C) Representative immunofluorescence images of TA skeletal muscle cross-sections of mice treated with recombinant SAA1 (0.3 *μ*g) or vehicle control at day 6 of treatment (laminin, white; Pax7, red; DAPI, blue). Yellow arrows indicate Pax7+ cells. Scale bar, 100*μ*m. (D) Quantification of myofiber cross-sectional area (CSA) of TA muscles of mice treated with recombinant SAA1 or vehicle control at day 6 of treatment. N=4. (E) Quantification of number of Pax7+ MuSC in TA skeletal muscle cross-sections of mice treated with recombinant SAA1 or vehicle control at day 6 of treatment. N=3. (F) Representative immunofluorescence images of primary mouse myotubes treated for 48 h with control medium, CM from parental KPC cells, or CM from SAA1 KD KPC cells. Myotubes were stained for MF20 (red) and nuclei with DAPI (blue) (left). Quantification of myotube area normalized to the number of nuclei is shown (right). Scale bar, 50 μm. N=3. (G) Tumor weight measured five weeks after orthotopic implantation of parental or SAA1 KD KPC cells. N=4-6. (H) Quantification of tumor free body weight of mice transplanted with KPC wt or SAA1 KD cells or sham-treated controls at 5 weeks post cell injection. N= 4-6. (I) Quantification of weight of tibialis anterior muscle (TA), normalized to body weight of mice transplanted with KPC wt or SAA1 KD cells or sham-treated controls at 5 weeks post cell injection (mg/g). N= 4-6. (J) Quantification of weight of soleus muscle, normalized to body weight of mice transplanted with KPC wt or SAA1 KD cells or sham-treated controls at 5 weeks post cell injection (mg/g). N= 4-6. (K) Longitudinal treadmill performance assessment over 5 weeks in mice transplanted with KPC wt or SAA1 KD cells or sham-treated controls. Quantification of running distance (meters) measured weekly and normalized to body weight at each time point. N=4-6. (L) Longitudinal wire hang test assay over 5 weeks in mice transplanted with KPC wt or SAA1 KD cells or sham-treated controls. Hanging time (seconds) was recorded weekly and normalized to body weight at each time point. N=4-6. (M) Kapaln Meier representing survival in PDAC and PDAC SAA1 KD animals. N=4-16. Data are presented as mean ± SEM. Statistical significance was determined using one-way ANOVA or two-way ANOVA. *p < 0.05; **p < 0.01; ***p < 0.001; ****p < 0.0001.

To investigate whether tumor-derived SAA1 contributes to cachexia, we reduced SAA1 expression in KPC cells using CRISPR-Cas9-mediated gene editing. Independent edited clones showed approximately 50% lower SAA1 expression without altered proliferation (**Suppl. Fig. 6C-D**). Conditioned media from SAA1-edited KPC cells induced substantially less myotube atrophy compared to parental KPC CM (**Figure 4F**), implicating SAA1 in the atrophic activity of the tumor secretome. We next assessed the *in vivo* contribution of tumor-derived SAA1 by orthotopic implantation of parental or SAA1-edited KPC cells. SAA1 reduction did not affect primary tumor growth or BW (**Figure 4G-H**), but preserved TA muscle mass compared with mice bearing parental tumors, whereas soleus muscle mass remained unaffected (**Figure 4I-J**). Mice bearing SAA1-edited KPC tumors also displayed improved treadmill endurance, wire hang assay performance, and forelimb grip strength (**Figure 4K-L** and **Suppl. Figure 6E**). Notably, SAA1 reduction prolonged survival without affecting primary tumor growth (**Figure 4M**). Thus, tumor-derived SAA1 contributes causally to skeletal muscle wasting and dysfunction, and its reduction uncouples host deterioration from primary tumor progression and prolongs survival.

### SAA1 signals through TLR4 to elicit distinct stress responses in myofibers and MuSC

To identify the receptor mediating SAA1 signaling in skeletal muscle, we examined known SAA1 receptors ^41^ in our snRNA-seq dataset. Among SAA1 receptors detected in skeletal muscle populations (*Ager, Cd36, P2rx7, Scarb1, Tlr2,* and *Tlr4*), only *Cd36* and *Tlr4* were significantly increased in cachectic muscles (**Suppl. Figure 7A**). *Cd36* upregulation was restricted to myonuclei, while *Tlr4* expression increased in both myonuclei and MuSC (**Suppl. Figure 7B-C**). CD36 inhibition with sulfosuccinimidyl oleate (SSO) ^42–44^ did not prevent KPC-CM- or recombinant SAA1-induced myotube atrophy (**Suppl. Figure 7D**). In contrast, TLR4 inhibition with TAK-242 ^45–47^ rescued SAA1- or KPC CM-induced myotube atrophy and prevented loss of Pax7+ MuSC (**Figure 5A-B**). Lentiviral shRNA-mediated Tlr4 knockdown (knockdown efficiency of > 90%) phenocopied the effects of TAK-242 in MuSC (**Suppl. Figure 7E**), establishing TLR4 as a critical mediator of SAA1-induced muscle dysfunction.

**Figure 5.**
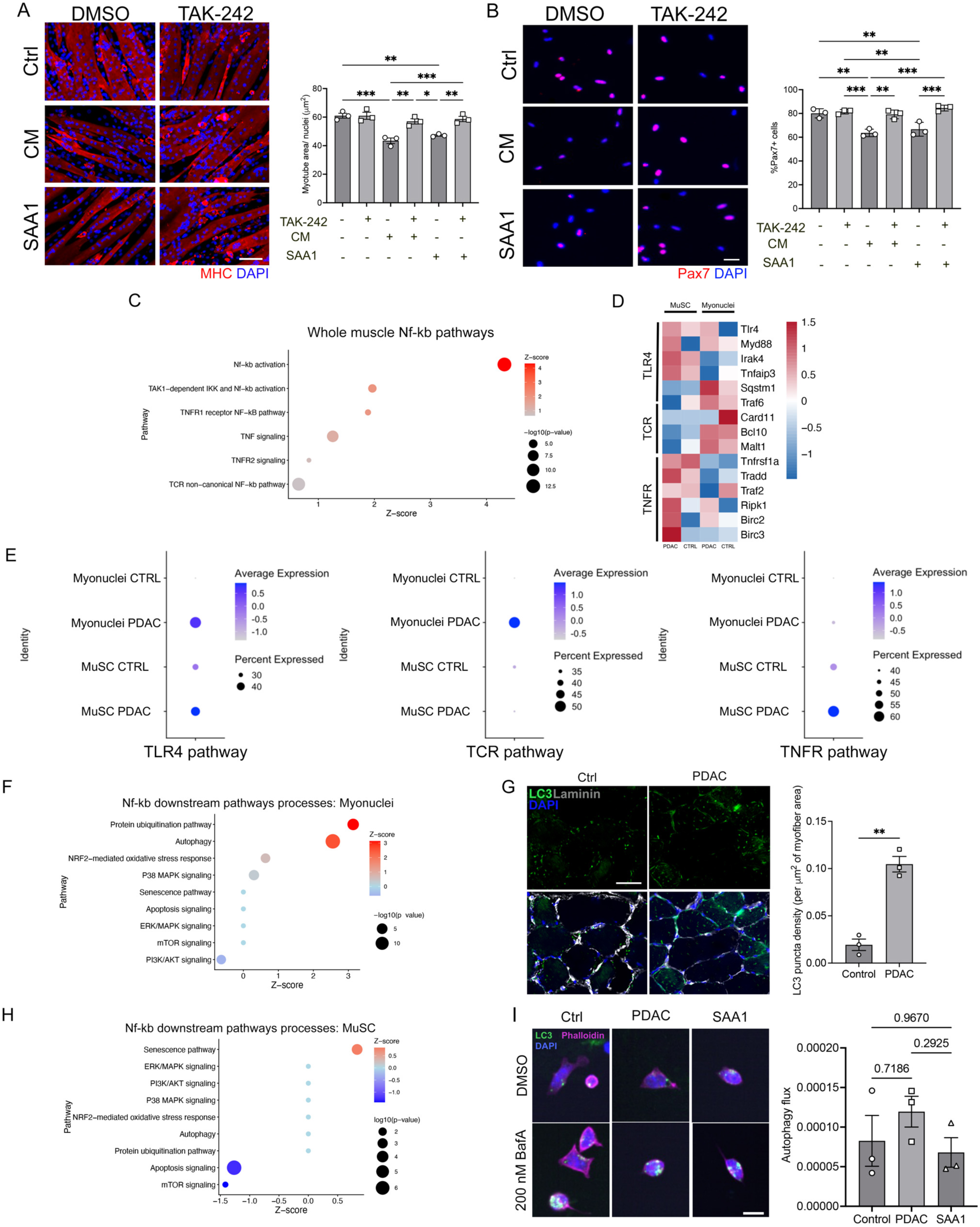
TLR4 signaling drives shared cachectic remodeling across skeletal muscle cell populations. (A) Representative immunofluorescence images of mouse MuSC-derived myotubes (left) treated with KPC conditioned media (CM), recombinant SAA1 (10 *μ*g/ ml), or vehicle control in the presence of absence of the TLR4 inhibitor TAK-242 (2 *μ*M) for 48 hours (myosin heavy chain, MF20, red; DAPI, blue). Scale bar, 50*μ*m. Quantification of myotube area normalized to the number of nuclei (right). N=3. (B) Representative immunofluorescence images (left) of mouse MuSC in growth conditions treated with KPC conditioned media (CM), recombinant SAA1, or vehicle control in the presence of absence of the TLR4 inhibitor TAK-242 for 48 hours (Pax7, red; DAPI, blue). Scale bar, 50*μ*m. Quantification of the percentage of Pax7+ cells in the indicated conditions (right). N=3. (C) Ingenuity Pathway Analysis (IPA) of whole-muscle transcriptomic data highlighting enrichment of NF-κB-related signaling pathways in cachexia. (D) Heatmap of NF-κB signaling modules across cell populations, subdivided into major upstream regulatory axes (TLR4-dependent, TNFR-mediated, and T cell receptor– associated signaling), in type IIb/atrophic myonuclei (right) and MuSC (left). (E) Module score analysis of NF-κB pathway activity (TLR4-, TCR-, and TNFR-associated signatures) in myonuclei and MuSC, confirming differential activation across cell types. (F) IPA of NF-κB downstream pathways in myonuclei (atrophic and IIb) showing enrichment of proteolytic and catabolic programs, including ubiquitin-mediated proteolysis and autophagy. (G) Representative immunofluorescence images of LC3 staining in tibialis anterior (TA) muscle cross-sections from sham-treated or KPC-injected mice (LC3, green; laminin, gray; DAPI, blue). Scale bar, 100 µm. Quantification of LC3 puncta density in the indicated conditions (right). (H) IPA of NF-κB downstream pathways in MuSC highlighting increased senescence-associated signaling, with no significant enrichment of autophagy-related pathways. (I) Autophagic flux analysis in MuSC isolated from sham-treated or KPC-injected mice and control treated with recombinant SAA1 (LC3, green; phalloidin, magenta; DAPI, blue). Quantification of LC3 accumulation upon bafilomycin A treatment is shown for the indicated conditions. N = 3. Scale bar, 10 µm. For bioinformatic analyses, statistical significance was determined using methods appropriate for each analysis, including differential expression testing with multiple testing correction (Benjamini–Hochberg adjusted p-values). Only results with adjusted p < 0.05 are shown. Statistical analyses were performed using unpaired two-tailed Student’s t-test or one-way ANOVA, as appropriate. Significance is indicated as follows: *p < 0.05, ***p < 0.005. Data are presented as mean ± SEM.

We next investigated how TLR4 signaling is resolved across skeletal muscle cell populations. NF-κB, a major downstream effector of TLR4 and a hallmark of cancer cachexia ^48,49^, was broadly activated in cachectic muscle (**Figure 5C**), but cell type-resolved analyses revealed distinct downstream programs. TLR4-associated programs increased in both myonuclei and MuSC, whereas TNFR-associated programs were preferentially enriched in MuSC and TCR-associated programs in myonuclei (**Figure 5D-E**), demonstrating cell-type specific responses.

These transcriptional differences were accompanied by distinct stress responses. Cachectic myonuclei showed activation of ubiquitin-proteasome, autophagy, NRF2-mediated oxidative stress response, and p38 MAPK signaling, with suppression of PI3K/AKT signaling (**Figure 5F**). Consistently, cachectic myofibers accumulated LC3+ and p62+ puncta *in vivo* (**Figure 5G** and **Suppl. Figure 7G**), suggesting altered proteostatic homeostasis and engagement of autophagy-associated pathways. MuSC instead exhibited senescence-associated programs without increased canonical autophagy (**Figure 5H**). Autophagic flux analysis using Bafilomycin A confirmed no significant increase in LC3 accumulation in MuSC isolated from cachectic mice or following SAA1 exposure (**Figure 5I**).

Additional pathway analyses and functional studies supported compartment-specific roles for stress-associated signaling (**Suppl. Figure 7F, H**). Exposure of MuSC to KPC CM or recombinant SAA1 increased the proportion of p62+ and A20+ cells without altering TRAF6 expression (**Suppl. Figure 7I-L**). Importantly, p62 accumulation occurred in the absence of detectable changes in autophagic flux (**Figure 5I** and **Suppl. Figure 7J**), indicating activation of stress-signaling pathways rather than canonical autophagy. Pharmacological inhibition of p62 with PTX80 exacerbated myotube injury, resulting in extensive myotube fragmentation and Caspase-3 activation (**Suppl. Figure 7M**), whereas p62 inhibition rescued the loss of Pax7+ MuSC induced by KPC CM or recombinant SAA1 (**Suppl. Figure 7N**). Thus, SAA1-TLR4 signaling disrupts both myofiber and MuSC homeostasis but engages distinct cell-state-specific stress responses, providing a mechanism through which a common tumor-derived signal coordinates multicellular muscle dysfunction.

### Therapeutic TLR4 inhibition rescues skeletal muscle dysfunction during PDAC cachexia

To determine whether TLR4 inhibition could therapeutically rescue established muscle dysfunction, we treated tumor-bearing mice with daily intraperitoneal (i.p.) injections of the TLR4 inhibitor TAK-242 beginning four weeks after orthotopic KPC implantation, when muscle atrophy and functional impairment were already evident (**Figure 6A**). TAK-242 treatment had no detectable effects in healthy control mice across all parameters examined (**Suppl. Figure 8A-I**). TAK-242 did not restore tumor-free body weight (BW) (**Figure 6B**), but rescued TA and gastrocnemius muscle mass normalized to BW (**Figure 6C-D**), with no effect on the soleus (**Figure 6E**). Importantly, primary tumor burden was unchanged (**Figure 6F**), indicating that therapeutic benefit reflects protection of host tissues rather than impaired tumor growth. TAK-242 also partially restored myofiber CSA and Pax7+ MuSC abundance (**Figure 6G-I** and **Suppl. Figure 8J**). Functionally, TAK-242 improved muscle performance in treadmill endurance, wire hang assays and four-limb grip strength relative to vehicle-treated controls (**Figure 6J-K** and **Suppl. Figure 8K**), demonstrating recovery of muscle function in addition to muscle mass.

**Figure 6.**
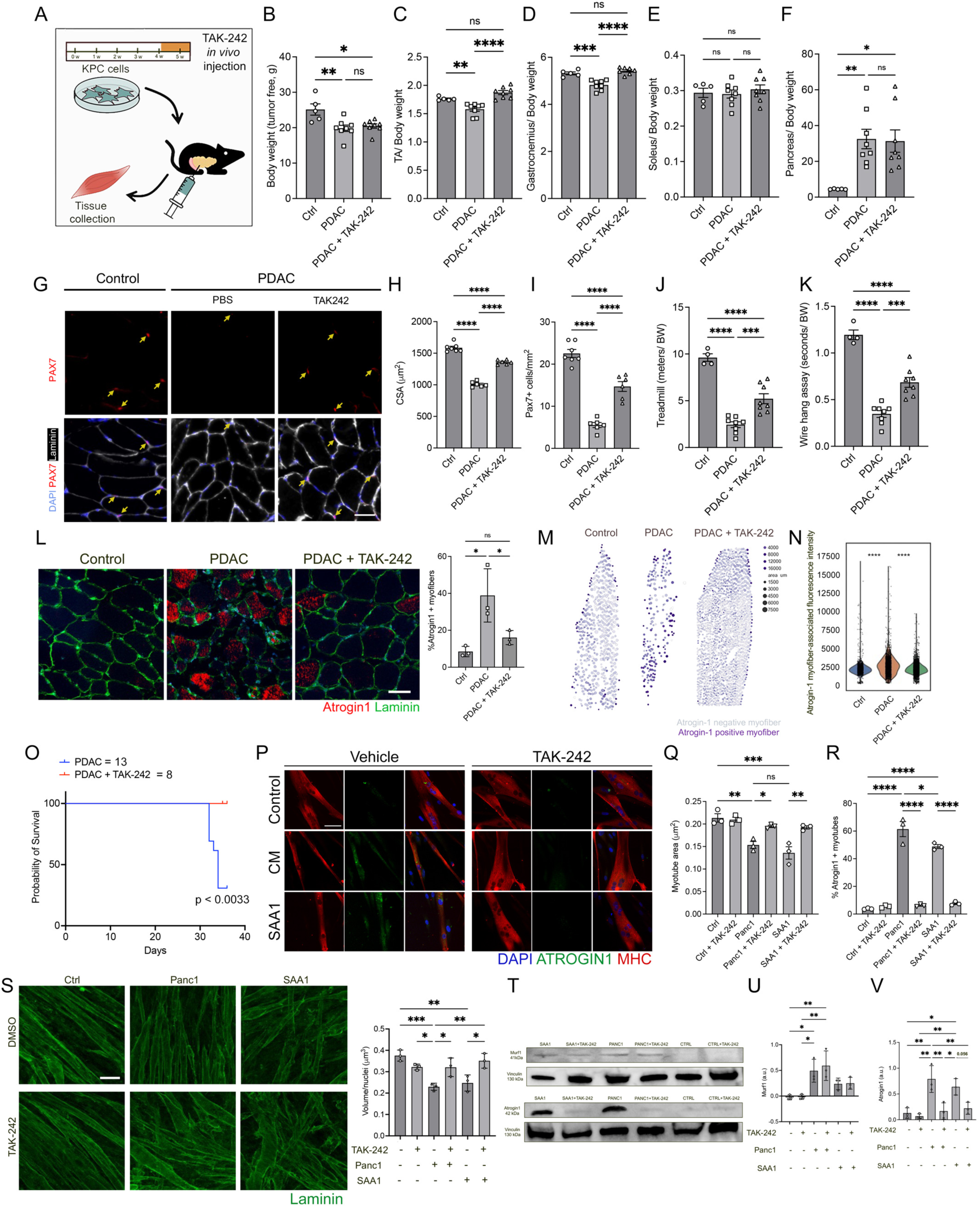
Pharmacological inhibition of TLR4 with TAK-242 mitigates cancer cachexia-induced muscle wasting, preserves MuSC content, and improves physical performance in PDAC-bearing mice and in human 3D engineered model. (A) Schematic of the *in vivo* treatment of PDAC or sham-treated controls with the TLR4 inhibitor TAK-242. (B) Quantification of tumor free body weight of mice transplanted with KPC cells or sham-treated controls with or without TAK-242 treatment (3 mg/kg) at 5 weeks post tumor implantation. N=5-8. (C) Quantification of weight of tibialis anterior muscle (TA), normalized to body weight of mice transplanted with KPC cells or sham-treated controls with or without TAK-242 treatment at 5 weeks post tumor implantation (mg/g). N=5-8. (D) Quantification of weight of gastrocnemius muscle normalized to body weight of mice transplanted with KPC cells or sham-treated controls with or without TAK-242 treatment at 5 weeks post tumor implantation (mg/g). N=5-8. (E) Quantification of weight of soleus muscles normalized to body weight of mice transplanted with KPC cells or sham-treated controls with or without TAK-242 treatment at 5 weeks post tumor implantation (mg/g). N=5-8. (F) Quantification of pancreas weight normalized to body weight of mice transplanted with KPC cells or sham-treated controls with or without TAK-242 treatment at 5 weeks post tumor implantation (mg/g). N=5-8. (G) Representative immunofluorescence images of TA skeletal muscle cross-sections of mice transplanted with KPC cells or sham-treated controls with or without TAK-242 treatment at 5 weeks post cell injection (laminin, white; Pax7, red; DAPI, blue). Scale bar, 100 µm. (H) Quantification of myofiber cross-sectional area (CSA) of TA muscles of mice transplanted with KPC cells or sham-treated controls with or without TAK-242 treatment at 5 weeks post cell injection. N=6-7. (I) Quantification of number of Pax7+ MuSC in TA skeletal muscle cross-sections of mice transplanted with KPC cells or sham-treated controls with or without TAK-242 treatment at 5 weeks post cell injection. N=6-7. (J) Treadmill performance assessment at week 5 in sham-treated controls or KPC-injected mice with or without TAK-242 treatment. Running distance (meters) was normalized to body weight at each time point. N = 4-8. (K) Wire hang test assay performed at week 5 in sham-treated controls or KPC-injected mice with or without TAK-242 treatment. Measurements were collected at the endpoint to compare treated versus untreated groups. Hanging time (seconds) was normalized to body weight at each time point. N = 4-8. (L) Representative staining by CODEX imaging of TA muscle cross-sections from sham-treated controls and KPC-injected mice with or without TAK-242 treatment (laminin, green; Atrogin-1, red) (left). Scale bar, 100 µm. Quantification of the percentage of Atrogin-1–positive myofibers across experimental groups (right). N= 3. (M) Atrogin1 spatial protein expression maps obtained by CODEX. Individual myofibers were computationally segmented and represented as dots preserving their spatial organization within the muscle cross-section. Dot color intensity reflects Atrogin-1 expression levels, highlighting the spatial distribution and heterogeneity of atrophic signaling across experimental groups. Dot size represents myofibers CSA. (N) Quantification of myofiber Atrogin-1 fluorescence intensity across sham-treated controls, KPC-injected mice with or without TAK-242 treatment. N=3. (O) Kaplan Meyer survival analysis of PDAC-bearing mice treated with vehicle (N = 13) or TAK-242 (N = 8). (P) Representative immunofluorescence images of human myotubes treated with Panc1 conditioned media (CM), recombinant SAA-1 (10 *μ*g/ml) or vehicle control, in the presence or absence of TAK-242 (2 *μ*M) for 48hrs. Myosin heavy chain (MF20, red), Atrogin-1 (green), nuclei (DAPI, blue). Scale bar, 50*μ*m. (Q) Quantification of myotube area of human myotubes treated with Panc1 conditioned media (CM), recombinant SAA-1 or vehicle control, in the presence or absence of TAK-242 (2 *μ*M) for 48hrs. N=3. (R) Quantification of the percentage of Atrogin-1+ human myotubes in cultures treated with Panc1 conditioned media (CM), recombinant SAA-1 or vehicle control, in the presence or absence of TAK-242 (2 *μ*M) for 48hrs. N=3. (S) Representative immunofluorescence images of 3D human myotube cultures treated with treated with Panc1 conditioned media (CM), recombinant SAA-1 or vehicle control, in the presence or absence of TAK-242 (2 *μ*M) for 48hrs. Laminin (green) (left). Scale bar, 50*μ*m. Quantification of myotube volume normalized to the number of nuclei in 3D human myotube cultures treated with Panc1 conditioned media (CM), recombinant SAA-1 or vehicle control, in the presence or absence of TAK-242 for 48hrs (right). N=3. (T) Representative Western blot for MuRF1, Atrogin-1 and Vinculin of protein lysates of 3D human myotube cultures treated with Panc1 conditioned media (CM), recombinant SAA-1 or vehicle control, in the presence or absence of TAK-242 for 48hrs. (U) Quantification of Western blot analysis for MuRF1 shown in (G) normalized to vinculin. N=3. (V) Quantification of Western blot analysis for Atrogin-1 shown in (G) normalized to vinculin. N=3. For CODEX analyses, only results with p < 0.05 are shown. Statistical analyses were performed using one-way ANOVA, Mann–Whitney U rank test or low-rank test as appropriate. Significance is indicated as follows: *p < 0.05, ***p < 0.005, ****p< 0.001. Data are presented as mean ± SEM. Statistical analyses were performed using one-way ANOVA followed by multiple comparisons testing as indicated in the figure legends. Significance is indicated as follows: *p < 0.05, **p < 0.01, ***p < 0.005, and ****p < 0.001. Data are presented as mean ± SEM.

Consistent with the structural and functional recovery, CODEX imaging showed reduced expression of Atrogin-1 and the stress-response factor ATF4 in TAK-242-treated myofibers (**Figure 6L-N** and **Suppl. Figure 8L-M**). Atrogin-1 induction was greatest in type IIb myofibers, with moderate increases in type IIx and IIa, and TAK-242 suppressed its expression across all affected fiber types (**Suppl. Figure 8L**). Notably, therapeutic TLR4 inhibition significantly prolonged survival without reducing primary tumor growth (**Figure 6O**). Thus, therapeutic TLR4 inhibition preserves muscle mass, restores muscle function and MuSC abundance, and improves survival independently of tumor growth, establishing the SAA1–TLR4 axis as a therapeutically actionable pathway in PDAC cachexia.

### The SAA1–TLR4 signaling axis is conserved in human skeletal muscle

Analysis of *rectus abdominis* muscle RNA-seq from patients with PDAC^11^ revealed increased *TLR4* expression in cachectic compared with non-cachectic muscle (**Suppl. Figure 9A**). To test the functional relevance of this pathway in human skeletal muscle, we exposed primary human myotubes to Panc1 CM or recombinant SAA1. Both treatments reduced myotube area and increased Atrogin-1, whereas TLR4 inhibition with TAK-242 rescued myotube size and suppressed Atrogin-1 induction (**Figure 6P-R**). TAK-242 similarly prevented the loss of Pax7+ human myogenic progenitors induced by Panc1 CM or SAA1 (**Suppl. Figure 9B**).

We further tested this pathway in 3D engineered skeletal muscle tissues ^50^ from primary human pericytes ^51,52^. Cells were differentiated within PEG-fibrinogen hydrogels under tension-driven conditions to generate aligned, contractile skeletal muscle tissues (**Suppl. Figure 9C**). Panc1 CM or SAA1 reduced myofiber volume normalized to nuclei number, which was rescued by TAK-242 (**Figure 6S** and **Suppl. Figure 9D-E**). Both treatments also increased Atrogin-1 protein, and this response was suppressed by TAK-242, whereas MuRF1 was not significantly altered (**Figure 6T-V**). Thus, SAA1–TLR4 signaling disrupts myofiber and myogenic progenitor homeostasis across complementary human skeletal muscle systems, supporting the translational relevance of targeting this pathway in cancer cachexia.

## Discussion

Our study identifies a previously unrecognized tumor-to-muscle signaling circuit through which pancreatic cancer disrupts skeletal muscle homeostasis. We show that tumor-derived SAA1 signals through TLR4 to promote myofiber atrophy while perturbing MuSC fate, and that targeting either the tumor-derived ligand or its host receptor preserves muscle function and improves survival without suppressing primary tumor growth. These findings establish the SAA1-TLR4 axis as a mechanism linking tumor-associated inflammation to host tissue dysfunction and expand the view of cancer cachexia beyond accelerated catabolism to a multicellular failure of tissue homeostasis.

Circulating SAA proteins are elevated in multiple malignancies and associated with poor outcomes ^53,54^, but their functional contribution to cachexia has remained unclear. Here, we demonstrate that SAA1 is highly expressed in PDAC tumor epithelial cells, secreted by pancreatic cancer cells, and persistently elevated throughout disease progression. Circulating SAA increased before overt muscle wasting and correlated with cachexia severity, while tumor SAA1 abundance was associated with lower BMI in patients. Importantly, partial genetic reduction of tumor-derived SAA1 attenuated muscle wasting and functional decline and prolonged survival without affecting tumor growth. Thus, SAA1 is not simply a marker of systemic inflammation but contributes directly to tumor-induced host deterioration. Its early and sustained elevation further raises the possibility that circulating SAA1 could help identify cachexia before substantial muscle loss, although prospective clinical studies will be required to establish its biomarker value. A central finding is that tumor-derived SAA1 disrupts skeletal muscle homeostasis at two complementary levels. SAA1 induced canonical atrophic remodeling in myofibers while promoting MuSC activation and myogenic commitment, consistent with progressive depletion of the MuSC pool. Collectively, these findings support a model in which cancer cachexia reflects not only accelerated tissue degradation but also disruption of the cellular mechanisms that maintain skeletal muscle homeostasis. Functional impairment emerged before detectable muscle atrophy, indicating that disruption of muscle homeostasis begins before substantial tissue loss. Together with our single-nucleus analyses showing remodeling across myogenic and stromal populations and reduced intercellular communication, these findings suggest that cachexia evolves through coordinated disruption of multiple cellular compartments rather than through myofiber catabolism alone. This framework may help explain why interventions focused solely on nutrition or anabolism have limited efficacy once tissue homeostasis is compromised.

Our findings also highlight context-dependent MuSC responses during cachexia. Previous studies in KPP and C26 models reported expansion of Pax7+ populations and impaired differentiation or fusion ^6,18^, whereas PDAC cachexia was characterized here by loss of MuSC quiescence, increased commitment and progressive depletion. Differences in tumor type, inflammatory milieu, and disease stage may therefore determine the nature of the stem cell response. More broadly, these observations identify maintenance of the regenerative compartment as an additional dimension of muscle preservation during chronic disease.

Our single-nucleus analyses further extend this framework by demonstrating that disruption of skeletal muscle homeostasis during cachexia extends well beyond the myofiber. We identified widespread remodeling of the muscle microenvironment, characterized by depletion of MuSC, FAP, and tenocytes, emergence of distinct atrophic type IIb myonuclear states, and profound disruption of intercellular communication networks. Trajectory analyses further revealed progressive activation of ubiquitin-proteasome and autophagy programs during the transition from healthy to cachectic myonuclear states, indicating that cachexia evolves through dynamic cell-state transitions rather than activation of a static atrophic program. Finally, the preferential vulnerability of glycolytic type II fibers, together with the relative resistance of soleus muscle, further supports the concept that intrinsic metabolic properties influence susceptibility to cachectic remodeling.

Mechanistically, our data identify TLR4 as a critical mediator of SAA1-induced muscle dysfunction. Pharmacological TLR4 inhibition and genetic suppression of TLR4 protected both myotubes and MuSC from SAA1- and tumor-CM-induced dysfunction. Despite this common upstream signal, myofibers and MuSC engaged distinct downstream stress programs. Cachectic myonuclei showed proteostatic, autophagic and oxidative stress responses, whereas MuSC exhibited senescence-associated programs without increased autophagic flux. The opposing effects of p62 inhibition in myotubes and MuSC further support cell-state-specific responses to inflammatory stress. Thus, a common tumor-derived signal can produce distinct pathological outcomes according to the identity and functional state of the responding muscle cell.

The therapeutic implications are supported by intervention after cachexia was established. TLR4 inhibition with TAK-242 restored muscle mass and partially recovered myofiber size, MuSC abundance, and muscle performance *in vivo* without reducing tumor burden, demonstrating that preservation of host tissue integrity can be achieved independently of direct anti-tumor activity. Notably, these improvements occurred despite failure to restore overall tumor-free body weight, suggesting that body weight alone may not capture meaningful recovery of tissue function. Both tumor SAA1 reduction and host TLR4 inhibition also prolonged survival independently of primary tumor growth, demonstrating that tumor progression and host deterioration can be mechanistically dissociated. These findings support therapeutic strategies directed at preserving host tissue function alongside tumor-directed treatment.

Several questions remain. SAA1 reduction produced substantial but incomplete protection, indicating that additional tumor-derived mediators contribute to PDAC cachexia. Whether SAA1-TLR4 signaling interacts with established cachectic cytokines or contributes similarly across tumor types remains to be determined. The effects of TLR4 inhibition should also be evaluated in combination with standard anticancer therapies, where preservation of muscle function could potentially improve treatment tolerance.

Together, our findings establish tumor-derived SAA1-TLR4 signaling as a mechanism linking pancreatic cancer to multicellular skeletal muscle dysfunction. By demonstrating that interruption of this signaling axis preserves muscle function and improves survival independently of primary tumor burden, our study identifies tumor-to-host communication as a therapeutically tractable component of cancer cachexia and provides a conceptual framework for interventions aimed at preserving host fitness during cancer treatment.

## Methods

### Contact for reagent and resource sharing

Further information and requests for resources and reagents should be directed to the Lead Contact, Alessandra Sacco, Ph.D.

### Experimental Animals

All protocols were approved by the Sanford Burnham Prebys Medical Discovery Institute Animal Care and Use Committee. Mice were housed according to institutional guidelines, in a controlled environment at a temperature of 22°C±1°C, under a 12-hour dark-light period and provided with standard chow diet and water ad libitum. Female C57BL6 mice at 8-10 weeks of age were used.

#### Orthotopic injection of PDAC KPC cells

For orthotopic tumor models, 2.5×10^4^ KPC.4662 cells were suspended in PBS with 50% matrigel (catalog number: CLS356231-1EA, Corning) and injected into the pancreata of C57BL6 mice, as previously reported ^26,27^. Cells were maintained under standard culture conditions and harvested at 70-80% confluency and then resuspended in PBS mixed 1:1 with ice-cold Matrigel, and injected in a total volume of 30 *μ*L. Mice were anesthetized with ketamine/xylazine (100/10 mg/kg provided by institutional animal facility) via intraperitoneal injection. Following sterile preparation, a left lateral abdominal incision was made to expose the pancreas. The cell suspension was injected into the tail of the pancreas using a 28G needle and the needle was held in place briefly to prevent leakage. The pancreas was returned to the abdominal cavity, and the peritoneum and skin were closed with sutures (peritoneum) and wound clips (skin). Mice received buprenorphine (3.25 mg/kg provided by institutional animal facility) subcutaneous for analgesia and were monitored until recovery and throughout the study. Wound clips were removed 9-10 days after the surgery.

At week 1, 2, 4, and 5 post-surgery, pancreas, tibialis anterior (TA), gastrocnemius, and soleus muscles were collected for downstream analyses. From day 30 to day 35 post-surgery, mice received daily intraperitoneal injections of TAK-252.

### *In vivo* and *in vitro* treatment with SAA-1 and TAK-242

SAA-1 recombinant protein (catalog number: #2948-SA R&D Systems, catalog number: TP302738 OriGene) was injected intramuscularly (10 *μ*g/ml in 30 *μ*l) into tibialis anterior muscles (TA) of mice. TLR4 inhibitor TAK-242 (catalog number: HY-11109 MedChem Express) was injected intraperitoneally (3 mg/kg in 100 *μ*l). Cells were treated *in vitro* with SAA-1 recombinant protein and TAK-242 for 48 h. SAA-1 was used at a final concentration of 10 *μ*g/ml, while TAK-242 was used at 1 μM. Both compounds were prepared according to the manufacturer’s instructions and diluted in culture medium immediately prior to treatment. DMSO or PBS-treated cells served as controls.

### Treadmill performance assay

Exercise capacity was assessed using a motorized treadmill (Columbus Instruments Exer-6M, serial 05664-2) equipped with an electrical stimulus grid at the rear. Mice were acclimated with two pre-training sessions performed during the 10 days prior to surgery, spaced 72 h apart, with the first session conducted the day before surgery. The test was performed once weekly. For testing, mice were placed on the treadmill at an initial speed of 5 m/min, with speed increased by 1 m/min every 30 s until exhaustion. Exhaustion was defined as the inability to disengage from the electrical stimulus grid for >20 s despite continued stimulation, at which point the test was terminated.

### Wire hang test assay

Muscle endurance and strength were assessed using a wire hang test. Mice were placed on a grid apparatus and allowed to grasp the surface with all limbs. Latency to fall was recorded as a measure of performance. Each mouse performed five trials, and the average latency was used for analysis. The test was performed once weekly.

### Cells

Mouse KPC cells derived from mice with the genotype Pdx1-Cre LSL-KRasG12D/+ LSL Tp53R172H/+ were provided by Cosimo Commisso and R. H. Vonderheide ^55^, and subcloned to give rise to the KPC#4662 clone used in this study. Human Panc1 cells were obtained from the American Type Culture Collection.

### Muscle Stem Cells Isolation

Muscle stem cells (MuSC) were isolated as described in Gromova et al., 2015 with minor revisions ^56^. Whole hindlimb (tibialis anterior, extensor digitorum longus, gastrocnemius, soleus, plantaris, vastus lateralis, quadriceps, and adductors) were minced and sequentially incubated in 800 units/mL collagenase type II solution (catalog number: 17101-015, Life technologies, Gibco^®^) and subsequent incubation with 80 units/mL collagenase II and 2 units/mL dispase II (catalog number: 04942078001, Roche) solution. Muscle tissues were then passed through a 10 mL syringe with 20 G needle and a 40 μm nylon filter. Primary and secondary antibodies incubation was performed in a 300 µl volume. Biotin-labeled lineage negative cells (CD45^+^, CD11b^+^, CD31^+^, Sca1^+^ cells) were either depleted using streptavidin beads (catalog number: 130-048-101, Miltenyi Biotec) through magnetic field or excluded during sorting by using streptavidin-APC-Cy7. MuSC were isolated with BD Biosciences FACSAria II cell sorter as CD45^−^, CD11b^−^, CD31^−^, Sca1^−^, CD34^+^ and integrin α-7^+^ population.

### Cell Culture

All cells were cultured in incubators at 37°C and 5% CO2. After isolation, murine MuSC were plated on laminin (catalog number: 11243217001, Roche, 1:25 dilution in PBS) in growth medium (catalog number: 10313-21, Gibco), 15% FBS (catalog number: FB-11, Omega Scientific), 1% Pen/Strep (catalog number: 15140163, Life technologies, Gibco^®^), 2.5 μg/mL FGFb (catalog number: 100-18B, Peprotech). For differentiation, cells were allowed to reach ∼90% confluence before switching to differentiation medium consisting of DMEM (Gibco), 1% Pen/Strep, and 2% horse serum (catalog number: 26050088, Thermo Fisher Scientific). Cells were maintained in differentiation conditions for 48–72 h to allow formation of mature myotubes. Subsequently, cells were treated with recombinant SAA1 (10 μg/mL) or KPC-conditioned media for 48 h. In each condition cells were further processed for immunostaining analysis. Murine KPC and Human Panc1 cells were cultured in growth medium composed of DMEM (Gibco) supplemented with 10% FBS and 1% penicillin–streptomycin. To generate conditioned media, cells were grown to 80–90% confluence, washed and incubated with fresh culture medium for 24 h. The conditioned medium was then collected, centrifuged at 500 x g for 5 min to remove cellular debris, and filtered through a 0.22 μm filter prior to use.

### *In vitro* treatment with PTX80 and Sulfosuccinimidyl oleate sodium

Cells were treated *in vitro* with sulfosuccinimidyl oleate (SSO; catalog number: HY-112847A MedChemExpress) and PTX80 (catalog number: HY-169779 MedChemExpress) for 48 h. SSO was used at a final concentration of 200 μM, while PTX80 was used at 1 μM. Both compounds were prepared according to the manufacturer’s instructions and diluted in culture medium immediately prior to treatment. DMSO-treated cells served as controls.

### Immunofluorescence

Muscle tissues (TA, gastrocnemius and soleus muscles) were isolated from healthy and PDAC-treated, SAA-1-treated or TAK-242-treated mice at different time points. Tissues were embedded in OCT and frozen in 2-methyl butane. Tissues were sectioned in 10 μm thick slices and further processed by immunostaining. Fixation was performed with 4% PFA (catalog number: sc-281692, Santa Cruz Biotechnology). Tissue sections were washed in PBS twice, then permeabilized with 0.5% Triton 100-X (catalog number: 1003477329, EMD Millipore) in PBS and blocked in 10% goat serum (catalog number: 16210-072, Life technologies, Gibco^®^) and 0.1% Triton 100-X in PBS at room temperature for 1 hour. For Pax7 staining, samples were incubated with AffiniPure Fab fragment goat anti-mouse IgG (1:40, catalog number: 115-007-003, Jackson ImmunoResearch) solution in 0.2 µm filtered PBS at room temperature for 30 mins, then washed with PBS, and incubated in antigen retrieval (1:100, antigen unmasking solution, citric acid based, catalog number: H-3300, Vector Laboratories) solution at 92°C for 10 mins. Incubation with the primary antibodies was conducted at room temperature for 1 hour or at 4°C overnight in blocking buffer. All washes after incubation with antibodies were done by using PBS with 0.5% Triton 100-X. For Pax7 staining (catalog number: Pax7-c, Developmental Studies Hybridoma Bank (DSHB), 1:10) in tissue sections, fixation was performed with 2% PFA for 20 mins at room temperature, permeabilization with -20°C methanol for 6 mins at room temperature, blocking and antibody dilutions in 5% BSA in PBS 1X, while washes were performed in 0.1% BSA in PBS 1X. Isolated cells were fixed in 4% PFA, washed in PBS twice, permeabilized at room temperature for 8 mins with 0.5% Triton 100-X, and incubated in 4% BSA (catalog number: SH30574.02, HyClone) with 0.5% Triton 100-X blocking buffer at room temperature for 1 hour or at 4°C overnight. The primary antibodies used are the following: mouse anti-Pax7 (catalog number: Pax7-c, Developmental Studies Hybridoma Bank (DSHB), 1:50 dilution for cultured cells, 1:10 for tissue sections and myofibers), mouse anti-myogenin (catalog number: 556358, BD Biosciences; 1:100 dilution), mouse anti-MHC (catalog number: Mf20-c, DSHB,1:50 dilution), rabbit anti-MyoD (catalog number: sc-760, Santa Cruz, 1:100 dilution), mouse anti-MyoD (catalog number: sc-377460, Santa Cruz, 1:100), rabbit anti-laminin (catalog number: L9393, Sigma, 1:100 dilution), rat anti-laminin (catalog number: 05-206, Millipore, 1:100 dilution), atrogin-1 (catalog number: AP2041, Antibodiesinc, 1:100 myotubes, 1:50 tissue dilution). Alexa-conjugated secondary antibodies (Invitrogen, 1:500 dilution) were diluted in appropriate blocking buffer depending on the type of sample and incubated at room temperature for 45-60 mins. Nuclear DNA was stained with DAPI (Catalog number: MBD0011, Sigma). Tissue section and myofibers were prepared for imaging in Fluoromount-G® (catalog number: 0100-01, SouthernBiotech) mounting solution. Images were acquired with Inverted IX81 Olympus Compound Fluorescence Microscope, XYZ Automated stage - ASI 2000 (Applied Scientific Instrumentation Inc.), with Color/monochrome cooled CCD camera - Spot RT3 and MetaMorph 7.11 Software (UIC, Molecular Devices) at 10X or 20X magnification or using confocal scanning through Leica TCS SP8 and LAS X software at 20X or 63X magnification. Leica DMi8 epifluorescent microscope was used for cell culture and tissue section imaging; a minimum of 8 fields of view acquired with 20X objective were analyzed. Nikon A1R HD confocal (running Nikon Elements software Version 5.42.04) with oil immersion 63X objective was used to acquire images of Pax7^+^ cells on myofibers. All images were composed, edited and modifications applied to the whole image using Adobe Photoshop 2025.

### Autophagic flux Assay *in vitro*

MuSC were isolated by FACS and seeded in a 384-well plate at a density of 1200-1300 cells in 100 µL MuSC media +/- 10 µg/mL recombinant SAA1 protein per well. Cells were incubated at 37°C, 5% CO2 for 24 hours. A media change was performed, replacing 50% of the MuSC media with 2X Bafilomycin A (AdipoGen #BVT0252M001) for a final concentration of 200 nM, or 2X DMSO, in MuSC media. Cells were incubated at 37°C, 5% CO2 for 2 hours, followed by fixation with 4% paraformaldehyde. Fixed cells were probed with 1:100 anti-LC3 (MBL #M152-3) in blocking buffer (0.2% saponin / 10% FBS/ DPBS), followed by 1:1,000 goat-anti-mouse Alexa 488 (Invitrogen # A11001) and 1X Alexa Fluor 647-labeled phalloidin (Thermo Scientific #A22287) in blocking buffer, and finally, 1:10,000 DAPI (Sigma #D9542) in DPBS. Imaging was performed on an Opera Phenix High-Content Screening system in confocal mode with a 40X objective. Total cell body and LC3-positive foci area was quantified by manually outlining and measuring the area of regions and of interest using a custom macro on ImageJ2 v2.16.0/1.54p. Autophagy flux was determined by subtracting the average total LC3-positive foci area per cell body area in DMSO-treated cells from that in Bafilomycin A-treated cells.

### Quantification of Muscle Tissue Cross-Sectional Area (CSA)

CSA quantification was performed in automated manner using a Macro through ImageJ64^57^ or Muscle Morphometry ImageJ plugin developed by Anthony Sinadinos (https://drive.google.com/drive/folders/0B_bBI7SbDQhCR1MxNEVXSlhiekE?resourcekey=0-8wdIKyTc0OKlB7WN67JqIw) by using the laminin fluorescent signal channel. CSA was calculated by using the laminin fluorescent signal. The area of each myofiber and their number in each field of view were obtained by converting the images into binary, then followed by the command “Analyze particles” limited to the set threshold value.

### snRNA sequencing

For each replicate (N=2 per condition), gastrocnemius muscle from 12-14-week-old mice was dissected, flash-frozen in liquid nitrogen, and stored at −80°C until processing. Muscles were thawed and minced in cold lysis buffer (10 mM Tris-HCl, 10 mM NaCl, 3 mM MgCl₂, 0.1% IGEPAL-CA630, 0.1% Tween-20, 1% BSA, and RNAse inhibitors: 0.15 U/µl Protector (Sigma, Cat#3335402001), 0.15 U/µl RNAseOUT (Thermo Fisher, Cat#10777019), 0.15 U/µl Superase-In (Invitrogen, Cat#AM2696), and 0.15 U/µl Watchmaker RNAse inhibitor (Watchmaker Genomics, Cat#7K0088-500UL) in DEPC-treated water). After mincing, 6 ml of cold lysis buffer was added and samples were incubated on ice for 10 min. Tissue was then homogenized using a Dounce homogenizer (8 strokes with a loose pestle), followed by dilution with 8 ml of wash buffer (10 mM Tris-HCl, 10 mM NaCl, 3 mM MgCl₂, 2% BSA, and the same RNAse inhibitor cocktail in DEPC water). The homogenate was sequentially filtered through 100 µm, 70 µm, and 40 µm cell strainers. Nuclei were pelleted by centrifugation at 500 × g for 5 min at 4°C using slow acceleration and deceleration settings and resuspended in 300 µl of cold resuspension buffer (1% BSA, 1× PBS, 0.2 U/µl RNAse inhibitor Protector). Nuclei were stained with 7-AAD for 15 min, followed by addition of 700 µl resuspension buffer and an additional centrifugation step under the same conditions. A final wash was performed with 500 µl resuspension buffer before resuspension in 300 µl for fluorescence-activated cell sorting (FACS). 7-AAD-positive nuclei were sorted using a BD FACSAria II into 500 µl collection buffer (2% BSA, 1× PBS, 0.4 U/µl RNAse inhibitor Protector), centrifuged at 350 × g at 4°C, and resuspended in 50 µl resuspension buffer. Nuclei were counted and adjusted to a final number of 20,000 nuclei per sample for downstream single-nucleus RNA-seq using the 10x Genomics Chromium 3’ v3 platform. Libraries were prepared according to the manufacturer’s protocol and sequenced on an Element Biosciences AVITI platform using paired-end 2 × 75 bp reads at a depth of ∼30,000 reads per nucleus.

### snRNAseq data processing and analysis

Raw sequencing data were aligned to the mouse reference genome (mm10-3.0.0 / GRCm38) using CellRanger (v8.0.1) with the --include-introns parameter. Ambient RNA was removed using CellBender (v0.3.0) ^58^, and doublets were identified with Scrublet (v0.2.3) ^59^ using an expected rate of 8%87. Downstream analyses were performed in R (v4.3.2) using Seurat (v5.2.1)88 ^60^. Across samples, an average of 16,438 nuclei and 988 genes per nucleus were recovered. After merging, nuclei were filtered to exclude those with <200 or >2,500 detected genes, mitochondrial content >5%, or Scrublet scores >0.2. Data were normalized using SCTransform, regressing out mitochondrial content, gene counts, cell cycle scores, and doublet scores. Batch correction across the three nuclei preparations was performed using Harmony integration (IntegrateLayers, SCT normalization, PCA reduction). Dimensionality reduction (PCA), clustering (FindNeighbors, FindClusters), and UMAP visualization (RunUMAP) were performed using the top principal components selected by elbow plot. Clustering was performed at resolution 0.2 (algorithm 3), and marker genes were identified using PrepSCTFindMarkers and FindAllMarkers. Major clusters were subset and reanalyzed to define subclusters, which were annotated based on differential gene expression, while contaminating populations were removed. Subclusters were then merged into a final integrated object and reprocessed as described above. Pathway enrichment analysis was performed using Ingenuity Pathway Analysis (IPA) on differentially expressed genes to identify significantly regulated biological processes. In parallel, pathway activity scores were computed using Seurat’s AddModuleScore function based on curated gene sets (>30 genes per pathway). Scores were calculated as the average expression of pathway-specific gene modules and compared across experimental groups. To characterize myonuclear dynamics, pseudotime analysis was performed using Monocle3 (v1.4.26) ^39,61,62^. The root node was defined based on canonical fiber-type identity for each analysis.

### RNA Isolation and Quantitative PCR

Total RNA was isolated with RNeasy Micro Kit (catalog number: 74004, Qiagen) following the manufacturer instruction. RNA quantification was performed with Qubit RNA HS Assay Kit (catalog number: Q32852, Invitrogen). The samples for qPCR analysis were further converted into cDNA with SuperScript^®^ VILO cDNA Synthesis Kit and Master Mix (catalog number: 11754050, Invitrogen) or High-Capacity cDNA Reverse Transcription Kit (catalog number: 4368814, Applied Biosystems) following manufacturer instructions. Real time PCR was performed on LightCycler^®^ 96 System (Roche) with Power SYBR**^®^** Green PCR Master Mix (catalog number: 4367659, Applied Biosystems), 5 µM primers concentration, and 0.5 ng of cDNA. Relative gene expression was calculated using the comparative Ct method, normalizing the Ct value of each target gene to Rpl13a or 18S rRNA (Rn18S). The primer sequences used were as follows: Atrogin-1/Fbxo32 forward, 5′-TGAGCGACCTCAGCAGTTAC-3′, reverse, 5′-TTCTCTTCTTGGCTGCGACG-3′; Rpl13a forward, 5′-AGCCTACCAGAAAGTTTGCTTAC-3′, reverse, 5′- GCTTCTTCTTCCGATAGTGCATC-3′; and 18S rRNA/Rn18S forward, 5′- GTAACCCGTTGAACCCCATT-3′, reverse, 5′-CCATCCAATCGGTAGTAGCG-3′.

### CRISPR-Cas9-mediated SAA1 gene editing

SAA1 editing KPC cell lines was achieved using a lentiviral CRISPR-Cas9 gene-editing approach. Single guide RNAs (sgRNAs) targeting the mouse Saa1 locus were designed and cloned into a lentiviral CRISPR-Cas9 expression vector, which was packaged into lentiviral particles using standard transient transfection protocols. KPC cells were transduced and selected with puromycin before undergoing single-cell cloning by limiting dilution. Individual clones were screened for SAA1 expression by ELISA of conditioned media through single cells plating, and genomic editing was confirmed by PCR amplification of the targeted genomic region followed by Sanger sequencing. Sequence chromatograms were analyzed using ICE (Inference of CRISPR Edits, Synthego) to estimate editing efficiency. Clones exhibiting an approximately 50% reduction in SAA1 secretion were selected for subsequent experiments.

### ELISA Assay

Cells were cultured for 48 hrs in growth conditions on laminin coated plates before medium collection. Upon collection, media was spun down for 15 s at highest speed (∼16,000 rpm) with a tabletop centrifuge. Supernatant was transferred into sterile tubes and stored at -80°C until analysis. Mouse SAA1-2 ELISA Kit (catalog number: KMA0021, ThermoFisher), Human SAA1-2 ELISA kit (catalog number KHA0011, ThermoFisher) and Human IL6 ELISA kit (catalog number ab178013) were used for this assay. Antibody cocktail (capture antibody, detector antibody, 5BR antibody - anti-SAA1-2) was prepared according to manufacturer directions. Undiluted sample, blank, and 6 serial dilutions of the sample were prepared and mixed with the antibody cocktail and then incubated in the provided plate for 90 minutes at room temperature on shaker at 400 rpm. After incubation, three 5-minute washes with wash buffer 1:20 were performed before 15 mins at room temperature incubation with TMB solution (in the dark). Immediately after, we added the stop solution and read at plate reader (FilterMax F5, Molecular Devices, operating on SoftMax Pro 7.0.2) at 450 nm. Quantification was performed using a standard curve generated from serial dilutions of the recombinant SAA standard, with background (blank) subtraction applied to all readings. Concentrations were interpolated by fitting a linear regression model in Microsoft Excel (Microsoft 365), deriving the equation (y = ax + b) and coefficient of determination (R² ≥ 0.967 across all assays) to assess fit quality. Sample values falling outside the linear range of the assay were excluded or re-analyzed within the dynamic range of the curve.

### Quantitative proteomics on serum using TMTpro labeling

Mouse plasma samples (2 µL) were solubilized using an equal volume of 2× SDS protein solubilization buffer containing 10% SDS and 100 mM triethylammonium bicarbonate (TEAB, pH 7.55). The proteins were reduced with 5 mM tris-(2-carboxyethyl)-phosphine for 30 min at room temperature and alkylated with 15 mM chloracetamide for 30 min at room temperature. Following acidification to a final concentration of 1.2% phosphoric acid, the samples were diluted with binding buffer (90% methanol, 100 mM TEAB, pH 7.1) and loaded onto S-Trap micro columns (Protifi). Proteins were digested on-column using sequencing-grade trypsin (Promega) at a 1:10 (w/w) ratio for 3 h at 47 °C. Peptides were eluted in a stepwise manner using 50 mM TEAB with 0.2% formic acid, followed by 50% acetonitrile with 0.2% formic acid, and then vacuum-dried. Peptide concentrations were measured via NanoDrop spectrophotometry, and 30 µg of total peptide was reconstituted in 100 mM HEPES (pH 8.5) for TMT labeling. TMTpro tags (ThermoFisher) in 100% anhydrous acetonitrile were added at a 3:1 (w/w) tag-to-peptide ratio to achieve a final peptide concentration of 1 µg/µL. The labeling reaction was incubated for 1 h at 25°C, quenched with 0.2% hydroxylamine for 15 min at 25 °C, pooled, and desalted using C18 TopTips (PolyLC) prior to drying. Dried peptides were reconstituted in 20 mM ammonium formate (pH ∼10) and fractionated on a Vanquish HPLC system using a Waters XBridge BEH C18 column (4.6 × 250 mm, 3.5 µm). Peptides were separated over a 45-min gradient up to 90% acetonitrile, and 48 collected fractions were non-contiguously pooled into 24 global proteome fractions and vacuum-dried. For LC-MS/MS, fractions were reconstituted in 2% acetonitrile with 0.1% formic acid and analyzed using a Proxeon EASY nanoLC coupled to an Orbitrap Fusion Lumos mass spectrometer equipped with a FAIMS Pro device (Thermo Fisher Scientific). Peptides were separated on a C18 Aurora column (75 µm × 250 mm, 1.6 µm; IonOpticks) at 300 nL/min using an 80-min gradient from 0% to 48% buffer B (80% acetonitrile, 0.1% formic acid). The mass spectrometer was operated in positive data-dependent acquisition mode, utilizing a three-experiment FAIMS method with compensation voltages of –45, –65, and –80 V and a 1 s cycle time per voltage. MS1 scans (m/z 350–1,500) were collected at 60,000 resolution with an AGC target of 4e5 and 50 ms maximum injection time. Precursor ions (charge states +2 to +7) were isolated with a 0.7 m/z quadrupole window and fragmented using higher-energy collisional dissociation (HCD) at a normalized collision energy of 35%. MS2 fragments were detected at 50,000 resolution with an AGC target of 5e4, an 86 ms maximum injection time, and a dynamic exclusion of 20 s. Mass spectra were processed using SpectroMine software (Biognosys, version 3.2.220222.52329) against the Uniprot mouse database. Searches required full tryptic specificity with a maximum of two missed cleavages. Carbamidomethylation of cysteine and TMTpro modifications on lysine and the peptide N-terminus were configured as fixed modifications, while methionine oxidation was set as a variable modification. A false identification rate of 1% was applied.

### CODEX spatial proteomics

PDAC tumors collected from orthotopic PDAC mice at 35 days post-injection, as well as tumors from TAK-242-treated and pancreas from sham control mice, were processed for CODEX imaging. Tissues were embedded in OCT, snap-frozen in liquid nitrogen–cooled isopentane, sectioned at 10 μm thickness, and mounted onto gelatine-coated coverslip. Sections were rehydrated in S1 buffer, fixed in 1.6% PFA for 10 minutes, blocked in S2 buffer for 5 minutes, and sequentially incubated overnight in a humidified chamber at 4°C with antibody cocktails in S2 buffer. Next, tissues were fixed in 1.6% PFA, further fixed using F fixative for 20 minutes, and stored in S4 solution prior to imaging as previously described ^63^. CODEX staining was performed using a multiplexed antibody panel including markers for immune, stromal, and muscle compartments, including MyoD, Pax7, MyHC isoforms, Laminin, Atrogin-1, and ATF4. Imaging was performed on a Phenocycler system (Akoya Biosciences) using a Keyence BZ-X810 automated microscope across 23 sequential cycles. Data acquisition and image reconstruction were performed using the CRISP Image Processor as previously described ^63^ and analyzed using python pipelines.

### Human subjects

Human skeletal muscle samples were obtained and processed as previously described^16^. Muscle biopsies from healthy donors’ patients were collected from lower extremity muscles during clinically indicated surgical procedures at Rady Children’s Hospital (San Diego), under Institutional Review Board approval and with written informed consent from legal guardians, in accordance with federal regulations for human subjects’ research. Human skeletal muscle-derived pericytes isolated as previously described ^64^ were cultured in complete medium (AmnioPrime, Capricorn Scientific GmbH, Germany) and embedded in PEG-Fibrinogen (PF) hydrogel (8 mg/mL fibrinogen) supplemented with 0.1% Irgacure 2959 photoinitiator in order to obtain 3D human muscle construct. Cells were suspended at 1 × 10⁴ cells/μL, and 80 μL of the cell-laden hydrogel was cast into Teflon molds equipped with U-shaped stainless-steel supports to promote tension-driven myotube alignment and differentiation ^64^. Constructs were polymerized under UV light (365 nm, 12 W, 5 min) and cultured in differentiation medium consisting of Amnioprime and DMEM (1:3) supplemented with 10% FBS.

### Human myogenic progenitor culture

Human skeletal muscle cells were plated on collagen (catalog number CLS354236-1EA, Millipore Sigma, 100 *μ*g/ml in acetic acid 0.02M) cultured in Dulbecco’s Modified Eagle Medium, 20% fetal bovine serum (FBS), and 1% penicillin–streptomycin. Culture medium was replaced every 48 h. For differentiation, cells were plated and grown to >85% confluence before switching to differentiation medium DMEM, 1% pen/strep, and 2% horse serum. Cells were maintained under differentiation conditions for 6 days to allow myotube formation, followed by treatment with Panc-1 conditioned media or recombinant SAA1 for 48 h.

### Analysis of human datasets (RNAseq and proteomics)

Human RNA-seq and proteomics datasets were retrieved from publicly available resources ^10,11,13,14^, and the DeepSpaceDB repository ^12,65^. Differential expression and pathway analyses were performed in R using standard statistical workflows. Ingenuity Pathway Analysis (IPA) was applied to the bulk RNAseq from Bailey et al. ^10^ and Bhatt et al. ^11^ datasets to identify enriched signaling pathways associated with cachexia-related transcriptional programs. The two patient cohorts were comparable in age, with median/mean ages of approximately 67 and 66 years, respectively. SAA1 expression was systematically evaluated across all datasets.

For the Cao et al. cohort, samples were stratified based on body mass index (BMI) into four groups (BMI >25, 22–25, 20–22, and <20). Expression values were normalized to the mean of the BMI >25 group, and relative expression changes were computed as ΔSAA1 (arbitrary units) by subtracting group-specific means from individual samples. This approach enabled cross-cohort comparison of SAA1 dynamics across metabolic states. For Bhatt et al., differential gene expression results were obtained directly from processed datasets provided by the senior author (Prof. Sambasivarao Damaraju) and integrated into downstream analyses.

### 3D hydrogel human myofibers

Three-dimensional constructs used in these experiments were generated using a synthetic hydrogel (PF) composed of polyethylene glycol (PEG) and fibrinogen (F) at a final concentration of 7.8 mg/mL. The mixture was supplemented with 0.1% of the photoinitiator Irgacure 2959 (1 mg/mL) to enable UV-induced polymerization. Cells were expanded to approximately 80% confluence, then detached, counted, and centrifuged at 1200 rpm for 8 minutes. Cell pellets were subsequently resuspended in PF at a final concentration of 1 × 10⁴ cells/μL. A total of 100 μL of bioink (containing 1 × 10⁶ cells) was then cast into Teflon molds equipped with a U-shaped stainless-steel support, designed to promote myogenic differentiation by applying mechanical tension and inducing parallel alignment of mature myotubes. Following casting, the cell-laden hydrogel was exposed to low-penetration UV light (365 nm, 12 W) for 5 minutes to allow complete polymerization. The resulting constructs were then transferred into culture plates containing AmnioPrime (Capricorn Scientific cat: #APR-B) medium to support cell growth and differentiation for 28 days.

### Western blot

Total protein extracts were obtained by lysing membrane fractions in a buffer containing 150 mM NaCl, 50 mM Tris–HCl (pH 7.5), 1% Nonidet P-40, 1 mM EGTA, 5 mM MgCl₂, and 0.1% SDS. The lysis buffer was supplemented with protease and phosphatase inhibitors, including 1 mM PMSF, 1 mM sodium orthovanadate, 1 mM NaF, and 1x protease and phosphatase inhibitor cocktails. Lysates were clarified by centrifugation at 14,000 rpm for 30 minutes, and protein concentration was quantified using a Bradford assay. Protein samples were denatured by incubation at 95°C for 10 minutes in NuPAGE LDS Sample Buffer supplemented with DTT as a reducing agent. Depending on experimental requirements, proteins were separated on 4-15% gradient polyacrylamide gels (Bio-Rad Mini-PROTEAN or Criterion systems). Following electrophoresis, proteins were transferred onto nitrocellulose membranes using the Trans-Blot Turbo transfer system (Bio-Rad). Membranes were blocked for 1 hour at room temperature in TBS containing 0.1% Tween-20 and 5% non-fat dry milk to reduce non-specific binding. Primary antibodies, diluted in blocking solution, were incubated with membranes overnight at 4°C. The following primary antibodies were used: rabbit anti-Murf1 (Prodotti Gianni 1739-T34), rabbit anti-Atrogin1 (Prodotti Gianni AP-2041) and mouse anti-vinculin (Ab-Cam AB18058). After washing, HRP-conjugated secondary antibodies were applied for detection. Signal was visualized using Clarity Western ECL substrate (Bio-Rad) and acquired with a LAS-3000 imaging system (Fujifilm). Band intensities were quantified using Fiji software and normalized to the appropriate loading control. All antibodies were used at dilutions recommended by the manufacturers.

### Statistics and Reproducibility

All statistical analyses were performed using GraphPad Prism version 7 (GraphPad Software) and R (version 4.x). Data are presented as mean ± standard error of the mean (SEM), unless otherwise indicated. The number of biological replicates (n) is specified in each figure legend. Normality was assessed using the Shapiro-Wilk test. Depending on distribution, statistical significance was evaluated using unpaired two-tailed Student’s t-test, one-way ANOVA, or two-way ANOVA for normally distributed data, and appropriate non-parametric tests for non-normal distributions. A p-value < 0.05 was considered statistically significant. Bioinformatic analyses were performed both on in-house generated datasets and publicly available transcriptomic and proteomic datasets ^10,11,13,14^ and DeepSpaceDB ^12^. Differential gene expression analyses, pathway enrichment (including Ingenuity Pathway Analysis), and comparative gene expression profiling were conducted using standardized R-based pipelines and/or processed datasets as described above. All statistical outputs derived from these analyses were considered significant only when meeting a threshold of p < 0.05 after the appropriate test and, when applicable, correction for multiple testing as implemented in the respective analytical frameworks. For the analysis of Supplementary Figure 1H using the Bhatt et al. dataset, expression levels of Trim63 and Fbxo32 were used to define cachexia status. Samples with expression above the cohort mean for both genes were classified as cachectic, whereas samples below the mean for both genes were defined as non-cachectic. This resulted in 80 cachectic and 4 non-cachectic samples. To account for extreme group imbalance, statistical comparisons were performed using a repeated random subsampling (Monte Carlo-based) approach, in which random subsets of four cachectic samples were iteratively sampled and compared against the non-cachectic group, generating a robust distribution of test statistics across iterations. Outlier samples were excluded based on predefined quality-control criteria, including abnormal distribution patterns, failure to cluster with biological replicates during unsupervised analyses, and/or evidence of technical variability. Outlier removal was performed prior to downstream statistical analyses to minimize technical confounding and ensure robust interpretation of biological signals. A minimum of three biological replicates was used for experimental validation, based on prior optimization within the laboratory to ensure statistical power for the assays performed. Randomization was applied at the level of group assignment, with mice randomly allocated to experimental conditions. Housing conditions were standardized and cage positions were rotated weekly to minimize environmental confounders. Blinding was implemented during data acquisition or analysis.

## RESOURCES AVAILABILITY

### Lead contact

Requests for further information and resources should be directed to and will be fulfilled by the lead contact, Alessandra Sacco.

### Materials availability

All unique reagents generated in this study are available from the lead contact with a completed material transfer agreement.

### Data and code availability

The snRNA-seq data generated in this study have been deposited in the NCBI Gene Expression Omnibus (GEO) database under accession number GSEXXXX.

## ACKNOWLEDGEMENTS

This work was supported by the US National Institute of Health (NIH) grants P30 CA030199, NIAMS R01 AR077448 and NIA P01 AG073084 to PDA, NIAMS R01 AR056712, NIAMS R01AR076247 and NIGMS GM134712 to PLP, R01 CA254806 to CC, California Institute for Regenerative Medicine CIRM-EDUC 12813 to GG, LC and CN, California Institute for Regenerative Medicine (CIRM) training grant TG2-01162 to D.S, French Muscular Dystrophy Association (AFM) postdoctoral grant to JM (AFM 28921), MDA Development Grant to LC (MDA 953791) and NIA P30 AG068635 pilot grant to CN. MGT was supported by California Institute for Regenerative Medicine CIRM-EDUC 12813. MGT was also supported by an American Society of Hematology (ASH) Hematology Inclusion Pathway (HIP) Graduate Student Award. Sanford Burnham Prebys Medical Discovery Institute Core Facilities are supported by NCI Cancer Center Support grant P30CA030199, and the following Shared Instrumentation Grants: S10-OD030285 (Nanostring GeoMX Digital Spatial Profiler, Genomics Core), S10-OD032325 (Cytek Aurora Full Spectrum Flow Cytometer, Flow Cytometry Core), S10-OD032408 (Zeiss LSM980 Airyscan 2, Confocal, Imaging Core). We thank Boster Bio for kindly providing the TLR4 antibody. We thank the members of the Puri and Colas labs, in particular Drs. M. Rossi for their immense support for histological analysis. We thank Dr. Denis Guttridge for critical discussion of the results. We thank Dr. Rouven Arnold and Ms. Ashley Neil for technical support. We thank the researchers who made their datasets publicly accessible, especially Drs. Damaraju and Baracos, which was implemented in this study. We thank the researchers who made their datasets publicly accessible, including Drs. Sean M. Grimmond and Peter Bailey for the PDAC transcriptomic dataset generated in “Genomic analyses identify molecular subtypes of pancreatic cancer” (GSE36924), as well as Drs. Hui Zhang and Liwei Cao for the proteogenomic PDAC datasets and analytical resources generated in “Proteogenomic characterization of pancreatic ductal adenocarcinoma.” (https://www.linkedomics.org/data_download/CPTAC-PDAC/). We thank Dr. Shinpei Kawaoka for the Human Spatial Dataset DeepSpaceDB ^12,65^ (https://genomics.virus.kyoto-u.ac.jp/deepspacedb/). We thank Dr. Karen Duong-Polk in the Commisso laboratory for teaching orthotopic injection surgeries. We thank the following people at the SBP Core Facilities for technical support: D. Sandoval, A. Vasquez, B. Charbono and A. Charbono from the Animal Facility; Y. Altman and B. Portillo from the Flow Cytometry Core Facility; R. Porritt for the Genomics Core Facility; L. Boyd from the Cell Imaging Facility; G. Garcia and M. Sevilla from the Histology Core Facility; S. Maurya and W. Huang from the Proteomics Core. We apologize to authors whose papers we could not cite due to space limitations.

## AUTHOR CONTRIBUTIONS

G.G. and A.S. designed the study, wrote the manuscript, and prepared the figures. G.G. performed experiments, bioinformatic analyses, and data interpretation. D.S. performed the initial experiments leading to the study conception. J.M. performed orthotopic KPC injections, isolated nuclei for single-nucleus RNA sequencing, and generated the Seurat objects from raw sequencing data. A.C. assisted with microscopy, qPCR, and western blot experiments. E.G. generated and analyzed the human 3D muscle cultures, including pericyte isolation, tissue engineering, immunostaining, and western blot analyses. B.S. performed CODEX experiments and analysis. J.R.B. assisted with orthotopic injections and muscle functional analyses. T.M. performed autophagy flux experiments. M.G.T. generated the KPC SAA1-edited cell line. R.M. analyzed human datasets. B.L., C.N., S.D., D.M., Y.Z., and P.G. assisted with computational, quantitative, proteomic, or technical analyses. C.S., L.C., A.R.C., A.V., C.K., P.A., W.W., C.G., P.L.P., and C.C. provided scientific input, technical expertise, and/or project supervision. A.S. supervised the study and acquired funding. All authors reviewed and edited the manuscript.

## SUPPLEMENTARY FIGURE LEGENDS

**Supplementary Figure 1 (related to Figure 1). Spatial and transcriptomic characterization of acute phase response genes in human PDAC and association with muscle atrophy.**

(A) Ingenuity Pathway Analysis (IPA) of bulk RNA-seq data from the Bailey et al PDAC tumor dataset ^10^ (n = 456 pancreatic ductal adenocarcinoma tumors) showing enrichment of inflammatory and acute phase response pathways in tumor tissue.

(B) IPA of bulk RNA-seq data from the Bhatt et al PDAC muscle dataset ^11^ (n = 84 patients) demonstrating upregulation of inflammatory pathways in rectus abdominis muscle from PDAC patients.

(C) Spatial transcriptomic visualization of SAA2 and CRP expression in tumor biopsies from patients with pancreatic ductal adenocarcinoma (PDAC) and non-tumoral controls with Pancreatic Intraductal Papillary-Mucinous Neoplasm, generated using DeepSpaceDB (Visium 10x platform) ^12^.

(D-E) Spatial transcriptomic analysis of SAA2 (D) and CRP (E) expression in tumor tissue from patients with PDAC (n = 10) and non-tumoral controls with benign Pancreatic Intraductal Papillary-Mucinous Neoplasm (n = 9), using the DeepSpaceDB (visium 10x) ^12^. Quantification of the fraction of SAA2/CRP-positive spatial spots through Seurat.

(F-G) Stratification of SAA2 (F) and CRP (G) expression by tumor stage (early vs. late) in the Bailey et al PDAC tumor dataset ^10^. Expression values are reported as log2(CPM) normalized to healthy pancreas from UCSC Xena. N=456. Control = 0.

(H-I) Stratification of IL6 (H) and TNF (I) expression by tumor stage (early vs. late) in the Bailey et al PDAC tumor dataset ^10^. Expression values are reported as log2(CPM) normalized to healthy pancreas from UCSC Xena. N=456. Control = 0.

(A) Dot plot analysis of single-cell RNA-seq dataset from human PDAC tumor tissue from Steele et al ^13^, showing cell type-specific expression of SAA1 and SAA2 across tumor and stromal populations.

(A) Violin plot showing SAA1 expression in rectus abdominis muscle from PDAC patients in the Bhatt et al dataset ^11^, stratified based on molecular atrophy status. Patients were classified as “atrophic” if both TRIM63 and FBXO32 expression levels were above the cohort mean, and “non-atrophic” if both were below the mean. N=37.

(A) Quantification of secreted IL6 (ng/mL) by ELISA in conditioned media from human pancreatic cancer cells (PANC-1), compared to control growth media condition. N=3. For bioinformatic analyses, statistical significance was determined using methods appropriate for each analysis, including differential expression testing with multiple testing correction (Benjamini-Hochberg adjusted p-values). Only results with adjusted p < 0.05 are shown. Statistical analyses were performed using unpaired two-tailed Student’s t-test or one-way ANOVA, as appropriate. For panel (K), statistical significance was determined using a permutation-based resampling approach, in which the control group (n = 4) was repeatedly compared to randomly sampled subsets of atrophic patients (n = 4 per iteration) to account for group size imbalance. Significance is indicated as follows: *p < 0.05, **p < 0.01, ***p < 0.005, and ****p < 0.001. Data are presented as mean ± SEM.

**Supplementary Figure 2 (related to Figure 1 and Suppl. Figure 1). Common inflammatory signaling programs between human tumor and skeletal muscle in PDAC cachexia and individual data points for Figure 1**.

(A) Individual data points corresponding to the quantification shown in Figure 1C.

(B) Individual data points corresponding to the quantification shown in Figure 1E.

(C) Individual data points corresponding to the quantification shown in Supplementary Figure 1F.

(D) Individual data points corresponding to the quantification shown in Supplementary Figure 1G.

(E) Individual data points corresponding to the quantification shown in Supplementary Figure 1H.

(F) Individual data points corresponding to the quantification shown in Supplementary Figure 1I.

(G) Individual data points corresponding to the quantification shown in Figure 1F.

Data are presented as mean ± SEM. Statistical analyses were performed as described in the main and supplementary figures.

**Supplementary Figure 3 (related to Figure 2). Longitudinal and spatial characterization of muscle wasting and MuSC dynamics in PDAC cachexia.**

(A-D) Longitudinal quantification of tissue weight normalized to body weight at weeks 1, 2, and 4 post tumor implantation in mice transplanted with KPC cells or sham-treated controls. Tissues analyzed include tumor/pancreas (A), tibialis anterior (TA) (B), gastrocnemius (C), and soleus (D) (mg/g).

(E) Representative hematoxylin and eosin (H&E) staining of TA muscle cross-sections from mice transplanted with KPC cells or sham-treated controls at 5 weeks post cell injection. Scale bar, 200 µm.

(F) Spatial mapping of myofiber types in TA muscle cross-sections obtained by CODEX imaging in control and PDAC-bearing mice. Individual myofibers were computationally segmented and represented as dots preserving their spatial organization. Dot color indicates fiber type (type IIa, blue; type IIb, orange; type IIx, green), and dot size corresponds to myofiber cross-sectional area.

(G) Quantification of myofiber type distribution in TA muscles based on CODEX analysis. N= 3.

(H) Representative immunofluorescence images of soleus muscle cross-sections from weeks 1 to 5 post tumor implantation (laminin, grey; Pax7, red, DAPI, blue). Scale bar, 100 µm.

(I) Quantification of myofiber cross-sectional area (CSA) in soleus muscles across time points. N= 3.

(J) Quantification of Pax7⁺ MuSC in soleus muscle cross-sections across time points. N= 3.

(K) Serum proteomic analysis of mice at 4 weeks post KPC tumor implantation or sham-treated controls showing expression levels of SAA1 and SAA2. Protein abundance is reported as intensity (arbitrary units). N=4-5.

(L) Quantification of MuSC abundance expressed as number of cells isolated by FACS per mg of tissue. Data are normalized to the mean of control samples at week 1. Statistical analyses were performed using one-way ANOVA or multiple unpaired two-tailed Student’s t-tests, as appropriate. Significance is indicated as follows: *p < 0.05, **p < 0.01, ***p < 0.005, and ****p < 0.001. Data are presented as mean ± SEM.

**Supplementary Figure 4 (related to Figure 2). Individual data points for *in vivo* and *in vitro* phenotypic analyses of KPC-tumor bearing mice.**

**(A)** Individual data points corresponding to the quantification shown in Figure 2K.

**(B)** Individual data points corresponding to the quantification shown in Figure 2M.

**(C)** Individual data points corresponding to the quantification shown in Figure 2N.

Data are presented as mean ± SEM. Statistical analyses were performed as described in the main and supplementary figures.

**Supplementary Figure 5 (related to Figure 3). Single-nucleus transcriptomic analysis reveals loss of MuSC quiescence and activation of muscle catabolic programs in cachexia.**

(A) Dot plot showing expression of representative marker genes used for cell-type annotation across identified meta-clusters in single-nucleus RNA-seq data.

(B) Top 50 downregulated genes in cachexia identified by pseudobulk differential expression (DEG) analysis across whole muscle from KPC-injected and sham-treated control mice at 5 weeks post tumor implantation.

(C) Ingenuity Pathway Analysis (IPA) of downregulated genes in cachexia showing the top 10 suppressed pathways.

(D-F) Module score analysis (AddModuleScore) across all cell clusters for ubiquitin (D), proteasome (E), and autophagy-mediated (F) pathways. # denotes cell clusters with significantly higher pathway scores in cachexia relative to controls; clusters increased in controls are not indicated.

(G-H) Cell-cell communication analysis using CellChat in control (G) and cachexia (H) conditions, focused on MuSC.

(I) Representative immunofluorescence images of FACS-isolated MuSC from healthy wild-type mice treated for 48h *in vitro* with KPC conditioned media or recombinant SAA-1 (10 *μ*g/ml) (Pax7, red; MyoD, green, DAPI, blue) (left). Scale bar, 50 *μ*m. Quantification of the percentage of Pax7+ and MyoD+ cells (right). N= 4.

(J) Representative immunofluorescence images of FACS-isolated myogenic cells treated for 48h with KPC conditioned media or recombinant SAA-1 (MyoD, green; Myogenin, red; DAPI, blue) (left). Quantification of the percentage of Myogenin+ cells (right). N= 3. Scale bar, 50 *μ*m.

(K) qPCR quantification of Atrogin-1 expression in skeletal muscle (TA) from SAA1-injected and PBS-injected muscle 6 days after the first injection. Normalization to Rpl13a. N= 3.

(L) qPCR quantification of Atrogin-1 expression in skeletal muscle (gastrocnemius) from sham-treated and PDAC mice 5 weeks after tumor implantation. Normalization to 18s rRNA. N= 3.

For bioinformatic analyses, statistical significance was determined using methods appropriate for each analysis, including differential expression testing with multiple testing correction (Benjamini–Hochberg adjusted p-values). Only results with adjusted p < 0.05 are shown.

**Supplementary Figure 6 (related to Figure 3 and Suppl. Figure 5). Cachexia remodels skeletal muscle intercellular communication, whereas tumor-derived SAA1 depletion preserves muscle function.**

(A-B) CellChat analysis of single-nucleus RNA-sequencing data showing differential numbers of interactions (A) and differential interaction strengths (B) between cachectic and control skeletal muscle. Red indicates increased and blue indicates decreased communication in cachexia relative to controls. Rows indicate signaling source cell populations and columns indicate target cell populations.

(C) ELISA quantification of SAA1 concentration in conditioned media collected from parental KPC cells, Rosa control KPC cells, and SAA1 KD KPC cells. N=3.

(D) Representative immunofluorescence images of parental and SAA1 KD KPC cells stained for Ki67 (green) with DAPI (blue) (left) and quantification (right). Scale bar, 50 μm. N=3.

(E) Longitudinal four-limb grip strength performance assessment over 5 weeks in mice transplanted with KPC wt cells, KPC SAA1 KD cells or sham-treated controls. Quantification of strength measured weekly. N=4-6.

Data are presented as mean ± SEM. Statistical significance was determined using one-way ANOVA or two-way ANOVA. *p < 0.05; **p < 0.01; ***p < 0.001; ****p < 0.0001.

**Supplementary Figure 7 (related to Figure 4). Cachexia/SAA1 drives TLR4-dependent signaling in cachectic muscle and promotes MuSC dysfunction.**

(A) Bar graph showing expression of SAA1 receptors identified by pseudobulk single-nucleus RNA-seq in skeletal muscle, comparing sham-treated (blue) and KPC-injected (red) conditions. Receptors include Ager, Cd36, P2rx7, Scarb1, Tlr2, and Tlr4.

(B-C) Cell-type-specific expression of Cd36 (B) and Tlr4 (C) in MuSC and myonuclei.

(D) Representative immunofluorescence images of murine MuSC-derived myotubes treated for 48h with KPC-conditioned media or recombinant SAA-1, in the presence or absence of the CD36 inhibitor SSO (sulfo-N-succinimidyl oleate) (MF20, red; DAPI, blue) (left). Quantification of myotube area normalized to the number of MF20+ nuclei (right). Scale bar 50 *μ*m. N= 3.

(E) Representative immunofluorescence images of lentiviral mediated shRNA Tlr4 knockdown in MuSC (mCherry reporter). Infected cells were treated for 48h with KPC-conditioned media or recombinant SAA-1 (Pax7, green; mCherry, red, DAPI, blue)(left). Quantification of Pax7+/mCherry+ cells is shown, normalized to control lentivirus (right). Scale bar 50 *μ*m. N=3.

(F) Ingenuity Pathway Analysis (IPA) of autophagy- and apoptosis-related pathways in myonuclei under cachectic conditions.

(G) Representative immunofluorescence images of skeletal muscle sections from sham-treated controls or KPC-injected mice (P62, red; laminin, gray; DAPI, blue)(left). Quantification of P62 density normalized to myofiber area (μm²) (right). Scale bar 100 *μ*m. N= 3.

(H) IPA analysis of autophagy- and apoptosis-associated pathways in MuSC.

(I) Representative immunofluorescence images of MuSC treated for 48h with KPC-conditioned media or recombinant SAA1 (A20, green; P62, gray; TRAF6, red; DAPI, blue). Scale bar 50 *μ*m. N= 3.

(J-L) Quantification of A20 (J), P62 (K), and TRAF6 (L) percentage of positive cells. N=3.

(M) Representative immunofluorescence images of murine myotubes treated with KPC-conditioned media or recombinant SAA1 in the presence of absence of the p62 antagonist PTX80 (30 nM) (MF20, red; Caspase-3, green; DAPI, blue) (left). Quantification of Caspase-3+ myotubes (right). Scale bar 50 *μ*m. N= 4.

(N) Representative immunofluorescence images of MuSC (Pax7, red; MyoD, green; DAPI, blue) treated with KPC-conditioned media or recombinant SAA1, in the presence or absence of the p62 antagonist PTX80 (30 nM) (left). Quantification of Pax7+ cell frequency is shown on the right. Scale bar 50 *μ*m. N= 4.

For bioinformatic analyses, statistical significance was determined using methods appropriate for each analysis, including differential expression testing with multiple testing correction (Benjamini–Hochberg adjusted p-values). Only results with adjusted p < 0.05 are shown. Data are presented as mean ± SEM. Statistical significance was determined using one-way ANOVA or two-way ANOVA. *p < 0.05; **p < 0.01; ***p < 0.001; ****p < 0.0001.

**Supplementary Figure 8 (related to Figure 5). TAK-242 does not show any effect in sham-treated control mice.**

(A-E) Evaluation of the effect of TAK-242 treatment (3 mg/kg) in sham-treated control mice. Quantification of pancreas-to-body weight ratio (A), body weight (B), tibialis anterior (TA) muscle weight normalized to body weight (C), gastrocnemius muscle weight normalized to body weight (D), and soleus muscle weight normalized to body weight (E) (mg/g). N= 3-5.

(F) Representative immunofluorescence images of TA skeletal muscle cross-sections from control mice treated with TAK-242 or vehicle (laminin, gray; DAPI, blue). Scale bar, 100 µm.

(G) Quantification of myofiber cross-sectional area (CSA) in TA muscles from control mice treated with TAK-242 or vehicle. N=3.

(H-I) Functional assessment of control mice treated with TAK-242 or vehicle during the final week of the experimental timeline (day 30-35). (H) Treadmill performance and (I) wire hang (hindlimb holding) test normalized to body weight. N=3-5.

(J) Quantification of MuSC abundance expressed as number of MuSC isolated by FACS per mg of tissue across experimental groups. N=3-5.

(K) Four-limb grip strength assessment at week 5 in sham-treated controls or KPC-injected mice with or without TAK-242 treatment. N = 3.

(L) Dot plot of Atrogin-1 expression across skeletal muscle fiber subtypes identified by CODEX analysis in sham-treated controls, KPC-injected mice with or without TAK-242 treatment. Fiber types are classified as type IIa, type IIb, and type IIx within each condition. Dot size signifies the percentage of Atrogin1-positive cells in each group, and color intensity (red gradient) indicates the average expression level of Atrogin-1. N=3.

(M) ATF4 spatial protein expression maps obtained by CODEX. Individual myofibers were computationally segmented and represented as dots preserving their spatial organization within the muscle cross-section. Dot color intensity reflects ATF4 expression levels, highlighting the spatial distribution and heterogeneity of atrophic signaling across experimental groups (left). Quantification of myofiber-associated ATF4 fluorescence intensity across control, PDAC, and PDAC + TAK-242-treated mice (right). N= 3. Statistical analyses were performed using unpaired two-tailed Student’s t-test, one-way ANOVA or Mann–Whitney U rank test, as appropriate. Significance is indicated as follows: *p < 0.05, ***p < 0.005. Data are presented as mean ± SEM.

**Supplementary Figure 9 (related to Figure 6). Human skeletal muscle cultures recapitulate TLR4-dependent cachectic remodeling.**

(A) Bulk RNA-seq analysis showing normalized TLR4 expression in rectus abdominis muscle from control and cachectic patients in the Bhatt et al dataset ^11^. N= 68.

(B) Representative immunofluorescence images of NCAM-positive human muscle progenitor cell-derived cultures treated with PANC-1 conditioned media (CM), recombinant SAA1, or vehicle control, in the presence or absence of TAK-242 for 48h (Pax7, red; DAPI, blue) (left). Scale bar, 50 µm. Quantification of the percentage of Pax7+ cells in NCAM-positive human muscle stem cell-derived cultures across treatment conditions (right). N= 3.

(C) Schematic depicting the process of generation of 3D human muscle cultures (left) and an example of 3D muscle (right).

(D) Representative immunofluorescence images of 3D human muscle treated with PANC-1 conditioned media (CM) or recombinant SAA1 for 48h (28 days after 3D muscle preparation) (myosin heavy chain, MF20, red; DAPI, blue). Scale bar, 50 µm.

(E) Representative high-resolution imaging of 3D human skeletal muscle constructs treated with PANC-1 CM or recombinant SAA1 (laminin, green; DAPI, blue). Scale bar, 100 µm.

Statistical analyses were performed using one-way ANOVA. Significance is indicated as follows: **p < 0.01, ***p < 0.005, ****p<0.001. Data are presented as mean ± SEM.

