## Supplementary Figures for "Tumor-derived SAA1-TLR4 signaling drives tumor-to-muscle communication in pancreatic cancer cachexia"

#### Supplementary figure 1

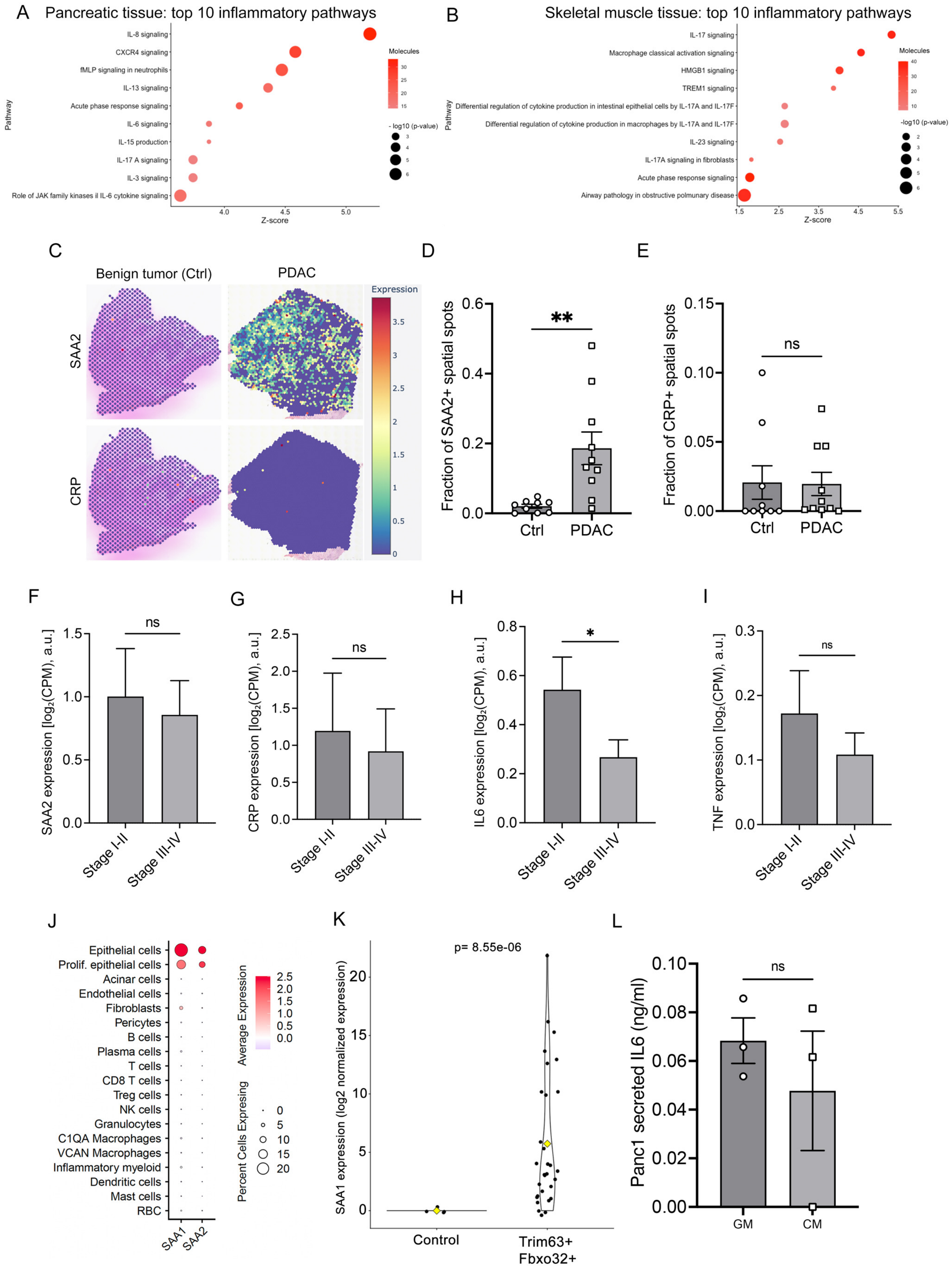

**Supplementary figure 2**

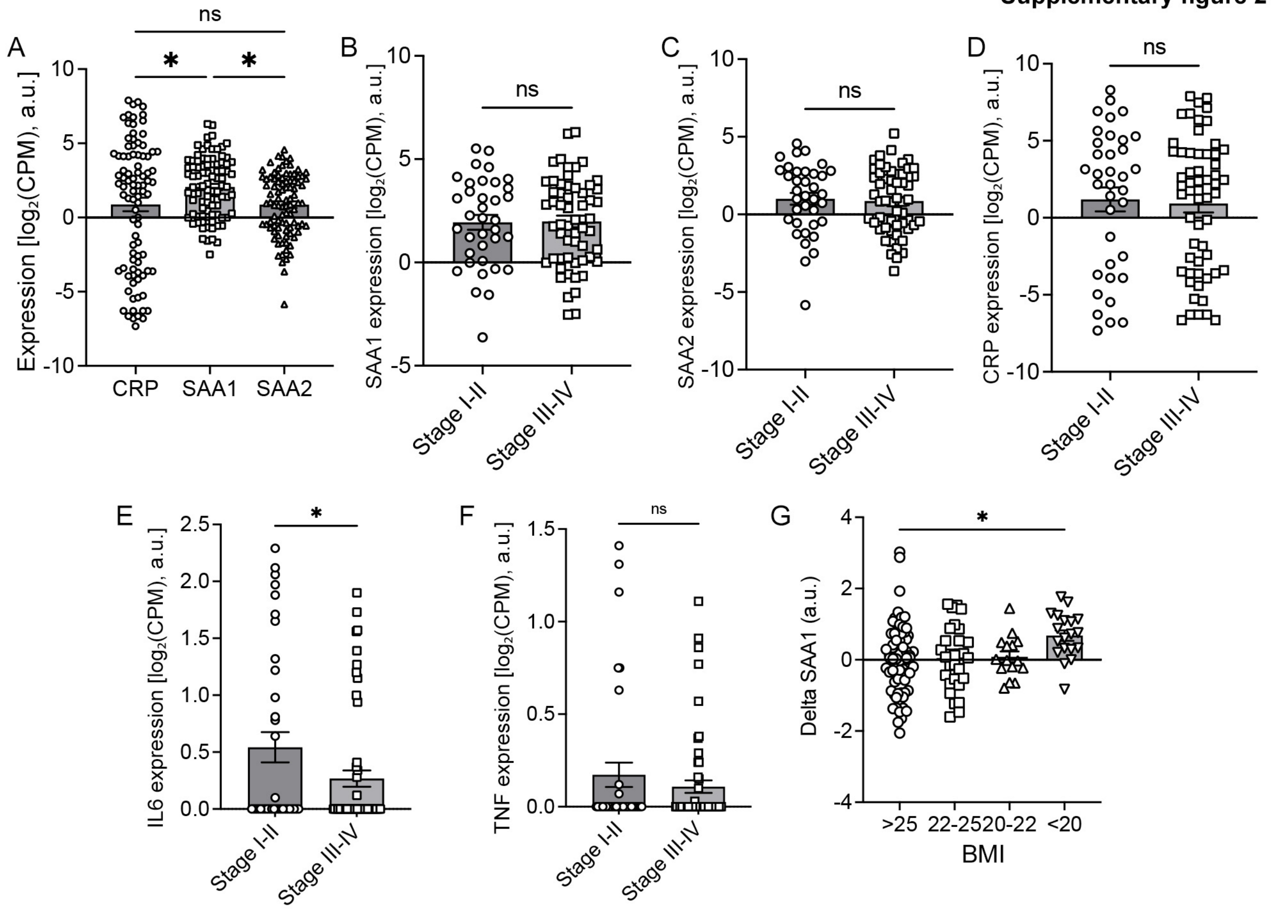

**Supplementary figure 3**

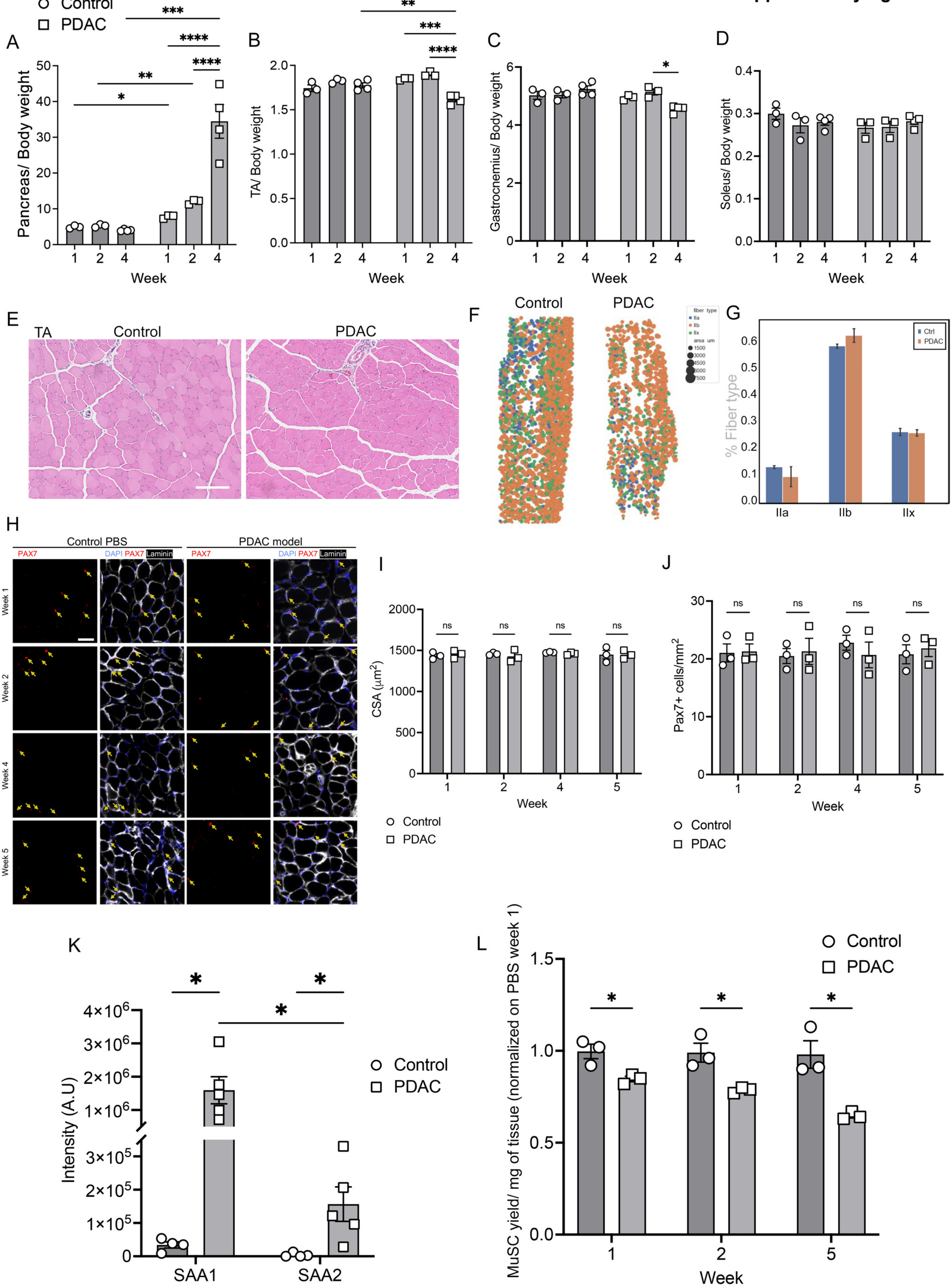

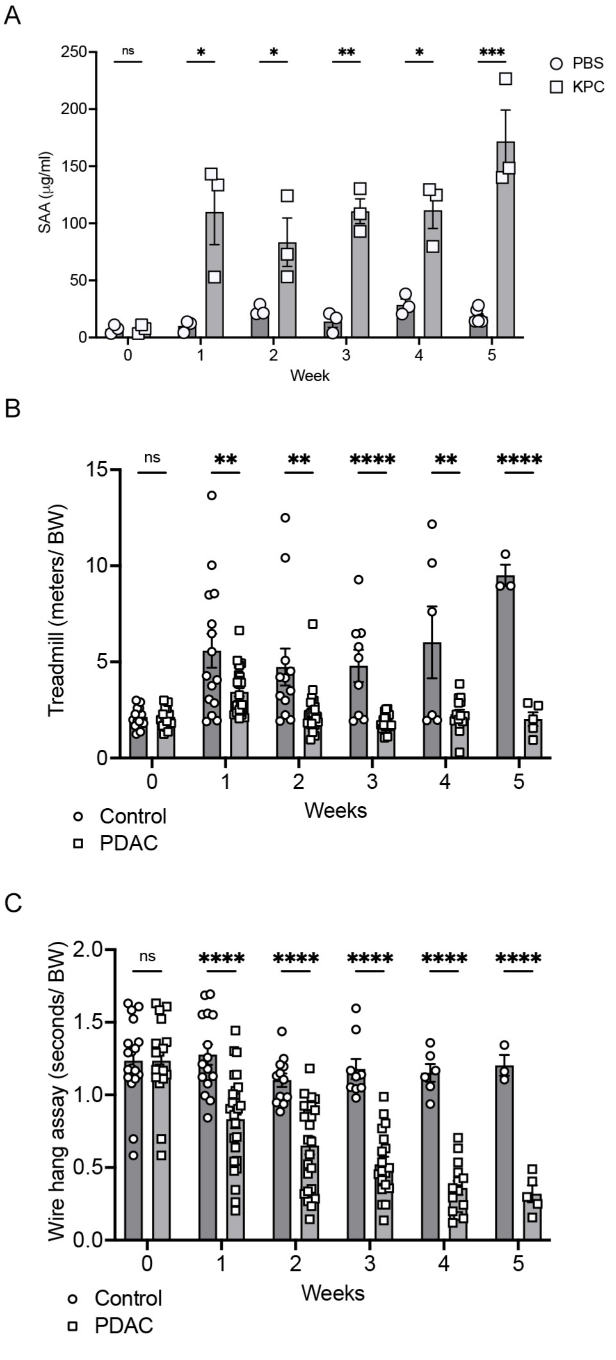

A
ns
*
*
**
*
***
PBS
KPC
SAA (µg/ml)
250
200
150
100
50
0
0
1
2
3
4
5
Week
B
ns
**
**
****
**
****
Treadmill (meters/ BW)
15
10
5
0
Weeks
Control
PDAC
C
ns
****
****
****
****
****
Wire hang assay (seconds/ BW)
2.0
1.5
1.0
0.5
0.0
Weeks
Control
PDAC

#### Supplementary figure 5

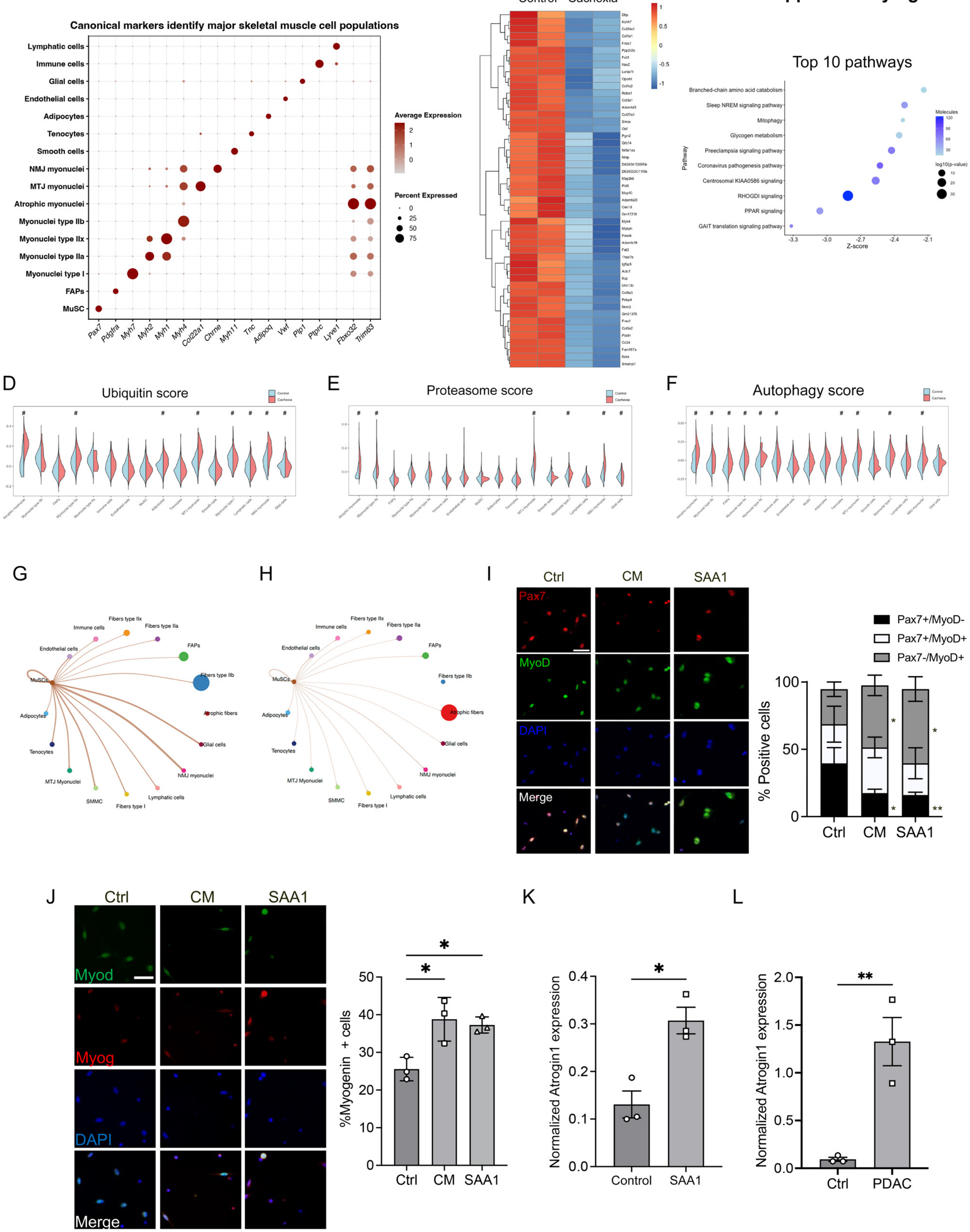

**Supplementary figure 6**

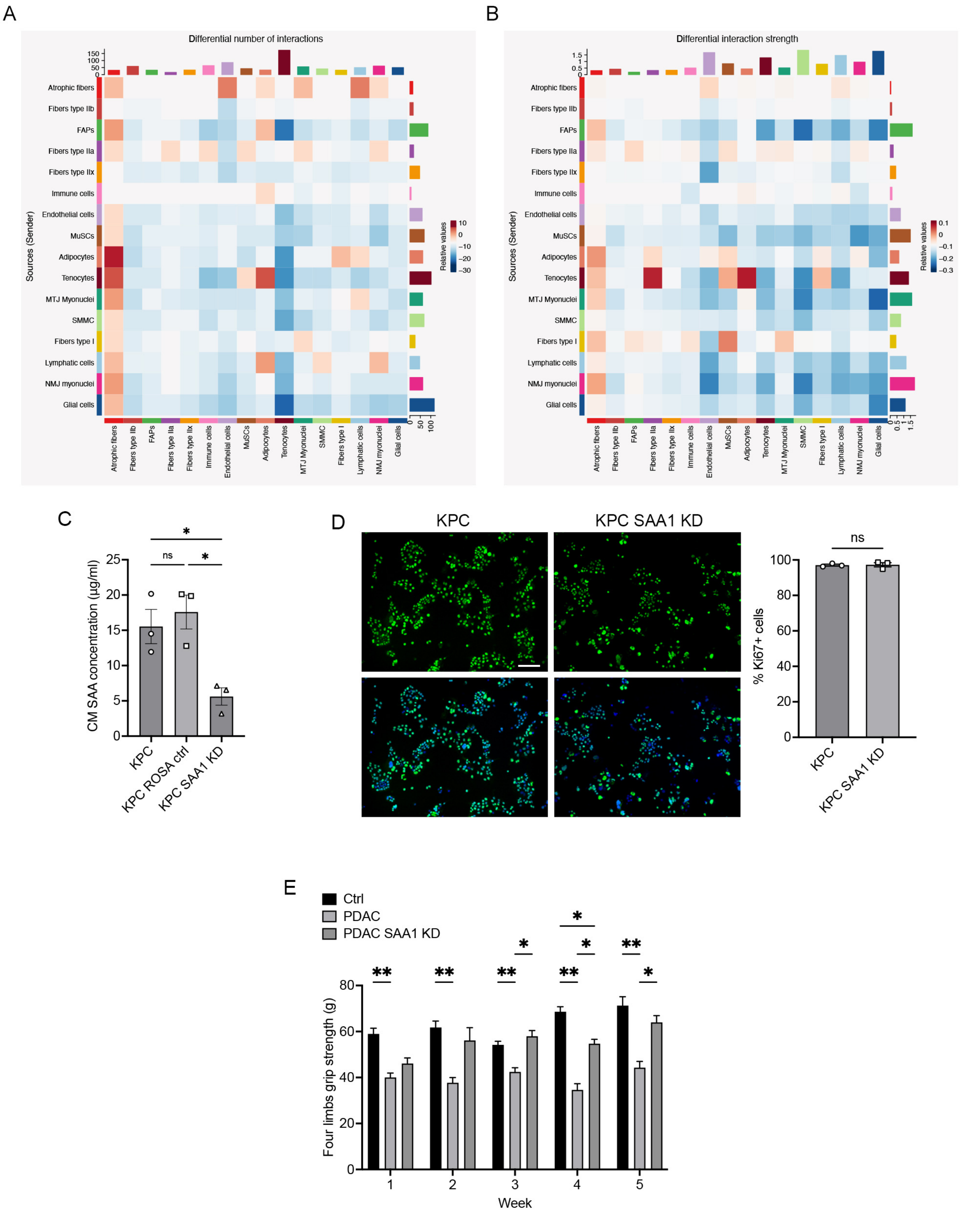

### Supplementary figure 7

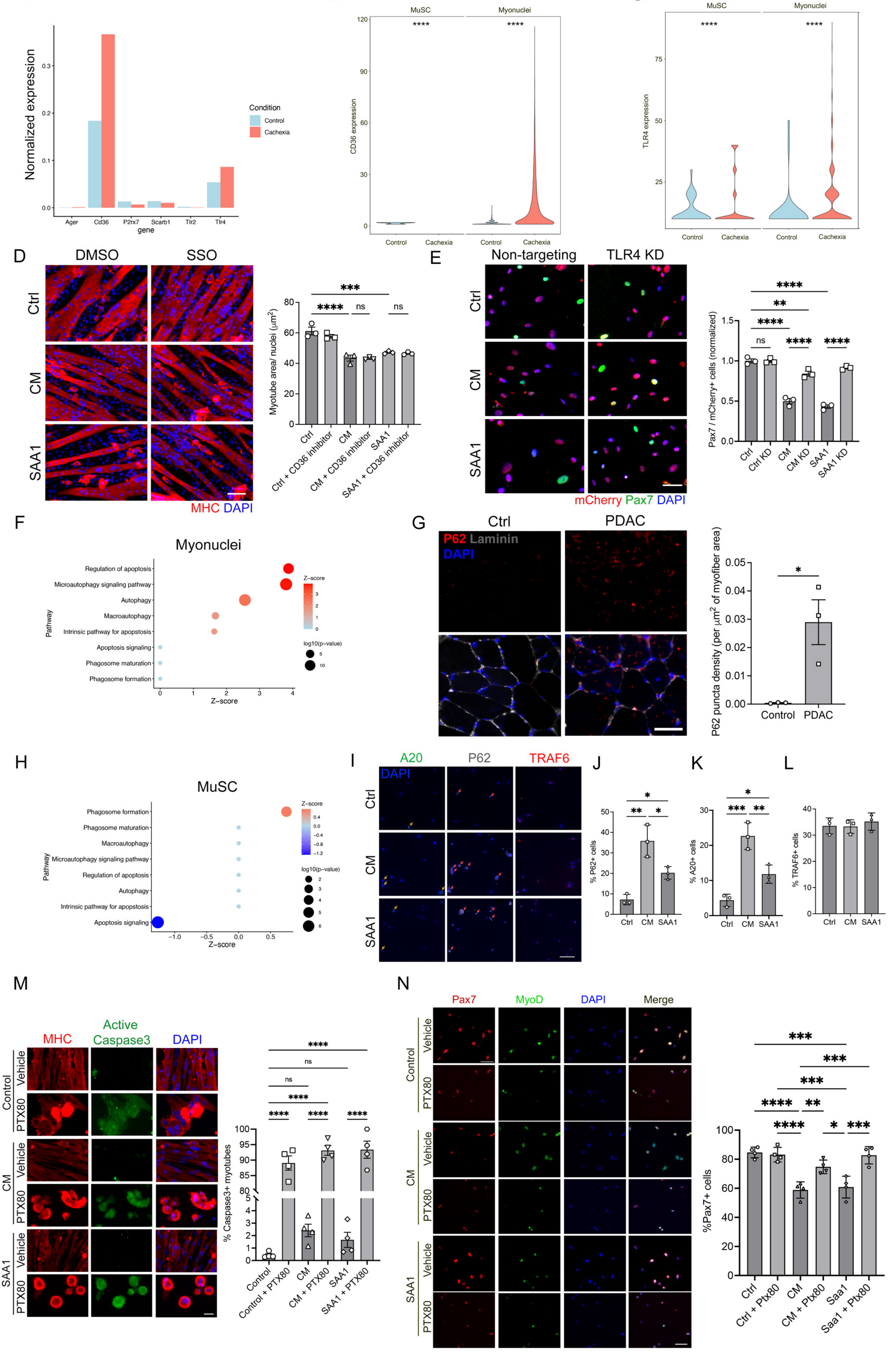

**Supplementary figure 8**

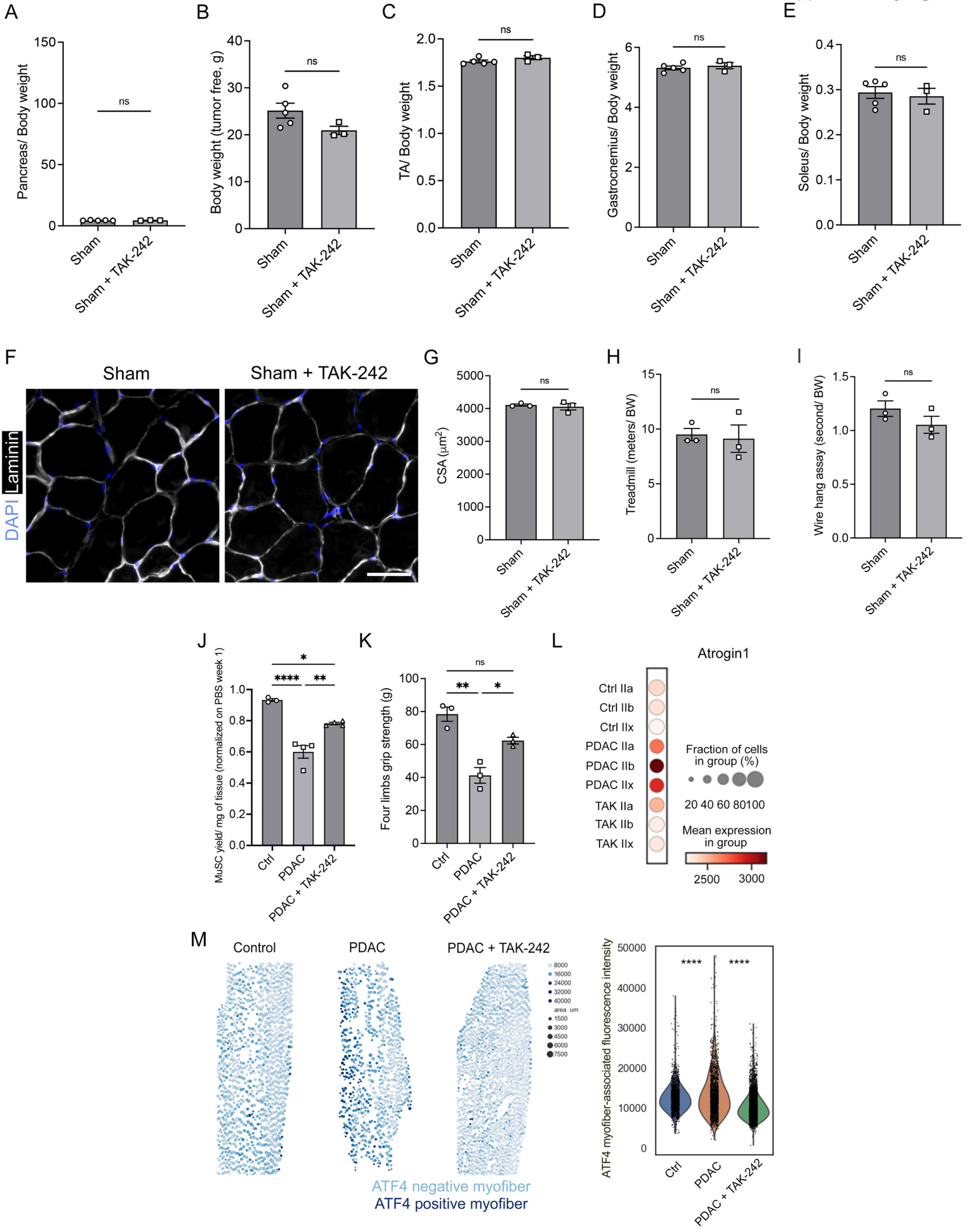

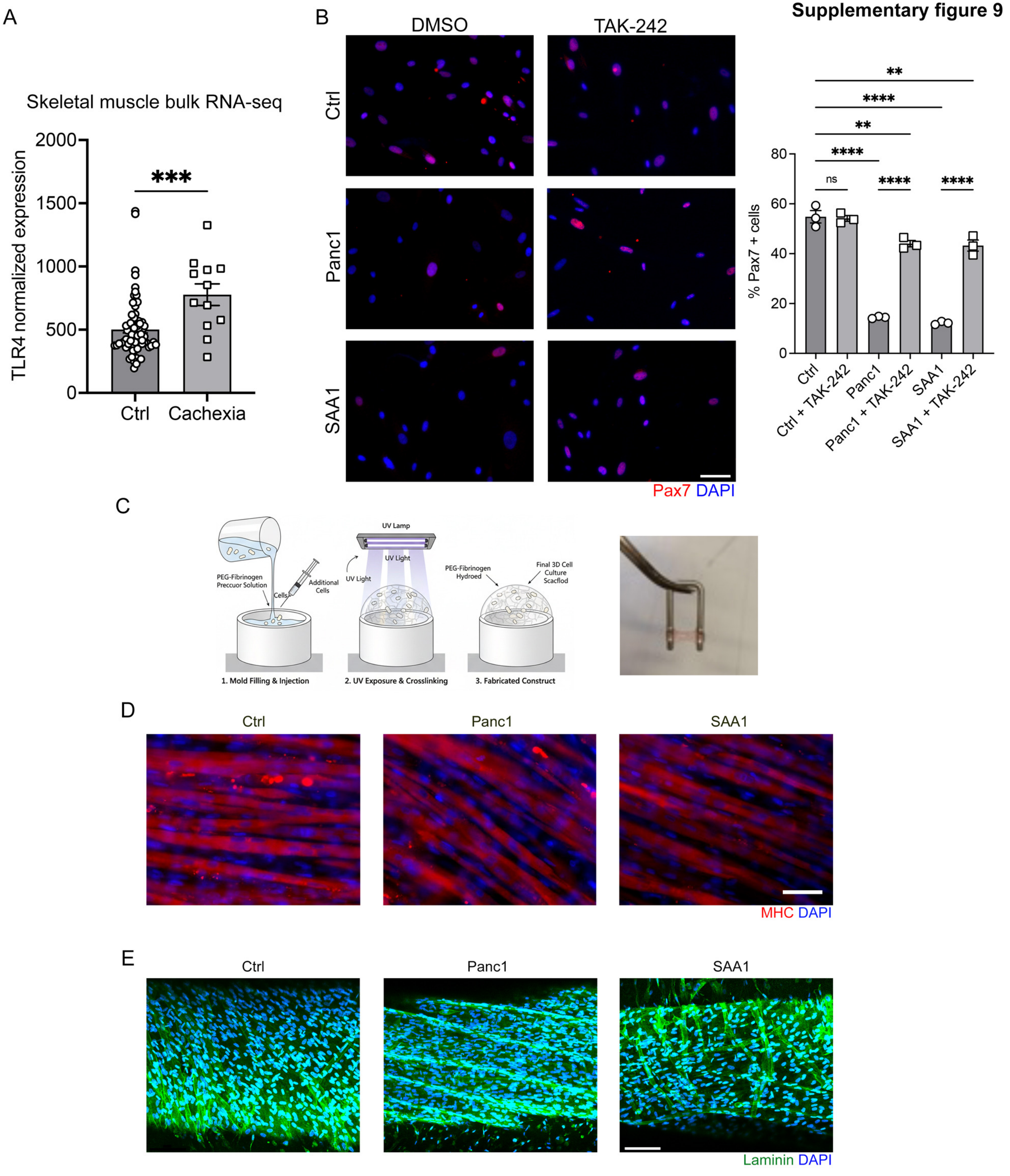
A
Skeletal muscle bulk RNA-seq
TLR4 normalized expression
2000
1500
1000
500
0
***
Ctrl
Cachexia
B
DMSO
TAK-242
Ctrl
Panc1
SAA1
Pax7 DAPI
% Pax7 + cells
80
60
40
20
0
ns
****
**
Ctrl
Ctrl + TAK-242
Panc1
Panc1 + TAK-242
SAA1
SAA1 + TAK-242
C
UV Lamp
UV Light
PEG-Fibrinogen Preccuor Solution
Cells
Additional Cells
PEG-Fibrinogen Hydroed
Final 3D Cell Culture Scacflod
1. Mold Filling & Injection
2. UV Exposure & Crosslinking
3. Fabricated Construct
D
Ctrl
Panc1
SAA1
MHC DAPI
E
Ctrl
Panc1
SAA1
Laminin DAPI
